# A peptide-centric quantitative proteomics dataset for the phenotypic assessment of Alzheimer’s disease

**DOI:** 10.1101/2022.11.04.515203

**Authors:** Gennifer E. Merrihew, Jea Park, Deanna Plubell, Brian C. Searle, C. Dirk Keene, Eric B. Larson, Randall Bateman, Richard J. Perrin, Jasmeer P. Chhatwal, Martin R. Farlow, Catriona A. McLean, Bernardino Ghetti, Kathy L. Newell, Matthew P. Frosch, Thomas J. Montine, Michael J. MacCoss

**Affiliations:** Department of Genome Sciences, University of Washington, Seattle, Washington 98195, United States; Department of Biomedical Informatics, Ohio State University, Columbus, Ohio 43210, United States; Department of Laboratory Medicine and Pathology, University of Washington, Seattle, Washington 98195, United States; Department of Medicine, University of Washington, Seattle, Washington 98195, United States; Department of Neurology, Washington University School of Medicine, 660 South Euclid Avenue, Box 8111, St. Louis, Missouri 63110, United States; Department of Pathology, Washington University School of Medicine, 660 South Euclid Avenue, Box 8111, St. Louis, Missouri 63110, United States; Massachusetts General Hospital, Department of Neurology, Harvard Medical School, 15 Parkman St, Suite 835, Boston Massachusetts 02114, United States; Department of Neurology, Indiana University School of Medicine, Indianapolis, Indiana 46202, United States; Department of Anatomical Pathology, Alfred Health, Melbourne VIC 3004, Australia; Department of Pathology and Laboratory Medicine, Indiana University School of Medicine, Indianapolis, Indiana 46202, United States; C.S. Kubik Laboratory for Neuropathology, and Massachusetts Alzheimer Disease Research Center, Massachusetts General Hospital, Boston, Massachusetts 02114, United States; Department of Pathology, Stanford University, Stanford, CA, 94305, United States

## Abstract

Alzheimer’s disease (AD) is a looming public health disaster with limited interventions. Alzheimer’s is a complex disease that can present with or without causative mutations and can be accompanied by a range of age-related comorbidities. This diverse presentation makes it difficult to study molecular changes specific to AD. To better understand the molecular signatures of disease we constructed a unique human brain sample cohort inclusive of autosomal dominant AD dementia (ADD), sporadic ADD, and those without dementia but with high AD histopathologic burden, and cognitively normal individuals with no/minimal AD histopathologic burden. All samples are clinically well characterized, and brain tissue was preserved postmortem by rapid autopsy. Samples from four brain regions were processed and analyzed by data-independent acquisition LC-MS/MS. Here we present a high-quality quantitative dataset at the peptide and protein level for each brain region. Multiple internal and external control strategies were included in this experiment to ensure data quality. All data are deposited in the ProteomeXchange repositories and available from each step of our processing.

## Background & Summary

Alzheimer’s disease (AD) is a major global public health problem. In the US, AD is the seventh most common cause of death for all ages and sexes. In contrast to ischemic heart disease, stroke and several forms of cancer, AD is increasing as a cause of death, of years lived with disability, and of disability-adjusted life years^1^. Success in limiting acute illnesses in the developing world is shifting the burden to non-communicable diseases, with an expected dramatic rise in AD globally by 2025^2^. Existence of forms of AD with known genetic causes or risk, and forms without known genetic underpinnings, highlight the potential for multiple molecular drivers and perhaps multiple pathogenic pathways involved in disease onset and progression. Moreover, longitudinal population-based cohort studies have repeatedly observed that AD is commonly comorbid with pathologic changes of vascular brain injury (VBI), Lewy body disease (LBD), limbic-associated TDP-43 encephalopathy (LATE), and/or hippocampal sclerosis^3,4^. AD is a chronic illness whose ultimate clinical expression as dementia follows years, if not decades, of injury, response to injury, consumption of reserve, and exhaustion of compensation. Determining the molecular profile of its various forms independent of comorbidities will be fundamental to efforts to develop tailored therapies that specifically target the molecular mechanism(s) of AD.

Thus, a primary focus of this data resource was the selection of tissue specimens based on current guidelines for AD neuropathologic change (ADNC)^3,4^ and specific exclusion for the presence of alternative potential causes of dementia, resulting in the carefully annotated examples of AD and controls free of medically significant comorbidities. Additionally, the specimens selected spanned the range of disease severity from the categories designated as high cognitive function (HCF) with no or low ADNC, HCF with intermediate or high ADNC, sporadic AD dementia (ADD, with intermediate or high ADNC), and autosomal dominant ADD (with causal mutations in *PSEN1, PSEN2*, or *APP*, and high ADNC). All research participants whose brains were used for this study underwent detailed, research quality, longitudinal cognitive assessments. For individuals without dementia, all had their last evaluation within 2 years of death and had neuropsychological test results in the upper quartile of the cohort to minimize interval conversion. All samples were obtained using a rapid autopsy protocol with postmortem interval less than 8 hours (except autosomal dominant AD due to practical limitations), flash frozen in liquid nitrogen, and kept frozen at -80°C prior to analysis. All samples were matched for age and sex except for those from individuals with autosomal dominant AD who experienced earlier onset.

Due to this careful specimen selection, we now have a unique sample set that can be used to study the molecular underpinnings of autosomal dominant ADD, sporadic ADD, and high burden ADNC with HCF without the confounding comorbidities faced in similar molecular profiling experiments. While previous studies have investigated the molecular profile of AD without excluding comorbidities, the high prevalence of these diseases, each of which can cause dementia on its own when present at high level, likely underlines the specificity of these profiles for AD. Likewise, because autosomal dominant AD is rare, typical molecular profiling studies have focused only on individuals with sporadic ADD, thereby limiting perspective on this heterogeneous disease. These points emphasize the uniqueness of this proteomics dataset for a more comprehensive assessment of the different forms of AD.

To make the most of this unique sample cohort, we used mass spectrometry proteomics methods that give reproducible and highly quantitative data. Most of the large-scale proteomics experiments studying the human brain have been performed using a mass spectrometry approach known as data-dependent acquisition (DDA). This approach is extremely powerful for building lists of proteins present in brain tissue and has been useful when combined with tandem mass tags for modest numbers of samples that can be performed within a “plex”. However, irregular sampling by DDA makes it challenging to provide robust and quantitative measurements across more samples than can fit in a plex. When using DDA, the number of peptides sampled is limited by the MS/MS sampling speed despite the dynamic range and peak capacity of the mass analyzer. A single MS spectrum can contain over one hundred different molecular species, of which only a handful are analyzed by MS/MS prior to the next full scan^5^. This general approach has become extremely powerful for cataloging proteins and modifications, but its irregular sampling results in missing data, requires extensive fractionation to sample low abundance peptides, and results in variable peptide sampling between runs of the same sample. Although the missing values in multiplexing tandem mass tags can be reduced at a protein level, it remains a problem on the peptide level due to the variable peptide sampling between runs^6^. In AD it is known that a subset of residues or peptides may be altered in disease, such as the Abeta region of amyloid precursor protein APP, or the proline-rich domain of Tau (MAPT)^2^. Consistent sampling and quantification on the peptide level is therefore necessary to accommodate such cases^7^.

An alternative to DDA is an acquisition approach known as data independent acquisition (DIA) that acquires comprehensive MS/MS information in a single LC-MS/MS run using a repeated cycle of wide-window MS/MS scans. The computational analysis of DIA spectra can be performed in the same “targeted” manner as fully targeted data; i.e., fragment ion chromatograms for each peptide can be extracted and used for quantification. However, unlike fully targeted data acquisition, DIA analysis can be done for any peptide in the sampled range (e.g., between 400 and 1000 *m*/*z*), rather than just for a subset of pre-specified peptides. Thus, the reproducible targeting and confident MS/MS-based quantification of parallel reaction monitoring (PRM) can be combined with DDA’s ability to detect and measure thousands of proteins. Like PRM methods, DIA requires reproducible chromatographic separation of peptides for reproducible quantification. This systematic sampling is important when scaling to data sets with samples prepared and run over many batches.

Typically, tens to hundreds of biological samples are processed and analyzed using LC-MS/MS in quantitative proteomics experiments. The regularity of DIA enables researchers to make peptide detections in one sample and use that information to inform the detection of the same peptides in other samples. DIA offers four key improvements over DDA. 1) Because peptides are sampled systematically, more peptides are detected in a DIA analysis than a DDA analysis in an equivalent length analysis^8,9^. 2) The same precursor *m*/*z* range is sampled, at the same RT, in all runs – eliminating the issues associated with stochastically sampled DDA data. 3) DIA analysis can make use of previously measured information to improve peptide measurement (e.g. known retention time, known fragmentation patterns, and which peptides provide stable and precise quantitative measurements)^10,11,12,13,14^. 4) Peptide detection can be assessed directly from DIA data, simplifying downstream analysis. These data provide an archive of all detectable molecular species within the measured mass range of the instrument. This methodology benefits from the reproducible and comprehensive sampling of the latest DIA methodology with an innovative approach used to improve peptide precursor selectivity^5^. The combination of the unique specimens with systematically collected mass spectrometry data creates a resource for the scientific community to test new hypotheses about the molecular features of different forms of AD dementia.

## Methods

### Human brain samples

Brain tissue samples were stratified into 4 groups based on clinical, pathological and genetic data and four brain regions (superior and middle temporal gyri or SMTG, hippocampus at the level of the lateral geniculate nucleus, inferior parietal lobule or IPL and caudate nucleus at the level of the anterior commissure). Cognitive status was determined as dementia or not dementia by DSM-IVR criteria. Individuals diagnosed as not dementia were from the Adult Changes in Thought (ACT) study and were included only if the last research evaluation was within 2 years of death and the last cognitive testing score using the cognitive abilities screening instrument (CASI) was in the upper quartile for the ACT cohort (>90); our definition of HCF. Brains from individuals with HCF who had no or low ADNC were designated “HCF/low ADNC” and those with intermediate or high ADNC were designated “HCF/high ADNC’’. All individuals diagnosed with ADD had intermediate or high level ADNC and were further subclassified as sporadic (“Sporadic ADD”) or ADD caused by a mutation in *PSEN1, PSEN2*, or *APP* (“Autosomal Dominant ADD”). Sporadic AD cases were from the ACT study and the University of Washington (UW) AD Research Center (ADRC), and Autosomal Dominant ADD cases were from the UW ADRC and the Dominantly Inherited Alzheimer Network (DIAN). Excluded was any case with LBD or LATE-NC other than involving amygdala, territorial infarcts, more than 2 cerebral microinfarcts, or hippocampal sclerosis. Time from death to cryopreservation of tissue, postmortem interval (PMI), was <8 hr in all cases except for those in the Autosomal Dominant ADD group. Details of sample stratification for the four brain regions (SMTG, Hippocampus, IPL and Caudate) are provided in Tables 1-4.

**Table 1.** Brain donor characteristics for the superior and middle temporal gyri (SMTG).

|  | HCF/low ADNC | HCF/high ADNC | Sporadic ADD | Autosomal Dominant ADD |
| --- | --- | --- | --- | --- |
| N | 9 | 11 | 18 | 24 |
| Age (*mean ± sd*) | 88 ± 5 | 90 ± 5 | 82 ± 13 | 51 ± 11 |
| Sex (*M:F*) | 4:5 | 5:6 | 11:7 | 15:9 |
| Post mortem interval hr (*mean ± sd*) | 3.9 ± 0.9 | 4.9 ± 1.5 | 4.5 ± 1.2 | 15.7 ± 9.3 |
| *APOE* ε4 alleles (*n (%)*) | 2 (11.1%) | 6 (27.3%) | 7 (19.4%) | 6 (17.6%)* |
| ε3:ε4 (*n*) | 2 | 4 | 5 | 4 |
| ε4:ε4 (*n*) | 0 | 1 | 1 | 1 |
| *missing* | 0 | 0 | 0 | 7 |
| Mutations (*n x gene*) | NA | NA | NA | 17 x *PSEN1*, 6 x *PSEN2*, 1 X *APP* |
| Braak Stage |  |  |  |  |
| B1 (*n*) | 4 | 0 | 0 | 0 |
| B2 (*n*) | 5 | 5 | 0 | 0 |
| B3 (*n*) | 0 | 6 | 18 | 24 |
| CERAD Score |  |  |  |  |
| C0 (*n*) | 9 | 0 | 0 | 0 |
| C2 (*n*) | 0 | 5 | 3 | 1 |
| C3 (*n*) | 0 | 6 | 15 | 23 |

\*For these; if there were missing genotypes the percentage is of the reported genotypes
Abbreviations: Alzheimer's disease (AD), AD dementia (ADD), AD neuropathologic change (ADNC), high cognitive function (HCF)

**Table 2.** Brain donor characteristics for the hippocampus.

|  | HCF/low<br>ADNC | HCF/high ADNC | Sporadic<br>ADD | Autosomal Dominant<br>ADD |
| --- | --- | --- | --- | --- |
| N | 10 | 11 | 20 | 3 |
| Age (*mean ± sd*) | 87 ± 6 | 90 ± 5 | 81 ± 12 | 53 ± 7 |
| Sex (*M:F*) | 5:5 | 5:6 | 12:8 | 2:1 |
| Post mortem interval<br>hr (*mean ± sd*) | 3.9 ± 0.9 | 5.1 ± 1.3 | 4.2 ± 1.4 | 8.5 ± 5.4 |
| APOE ε4 alleles (*n (%)*) | 2 (10.1%) | 6 (27.3%) | 10 (25.0%) | 2 (33.3%) |
| ε3:ε4 (*n*) | 2 | 4 | 6 | 0 |
| ε4:ε4 (*n*) | 0 | 1 | 2 | 1 |
| *missing* | 0 | 0 | 0 | 0 |
| Mutations (*n x gene*) | NA | NA | NA | 2 x *PSEN1*, 1 X *APP* |
| Braak Stage |  |  |  |  |
| B1 (*n*) | 5 | 0 | 0 | 0 |
| B2 (*n*) | 5 | 5 | 0 | 0 |
| B3 (*n*) | 0 | 6 | 20 | 3 |
| CERAD Score |  |  |  |  |
| C0 (*n*) | 10 | 0 | 0 | 0 |
| C2 (*n*) | 0 | 5 | 4 | 0 |
| C3 (*n*) | 0 | 6 | 16 | 3 |

**Table 3.** Brain donor characteristics for the inferior parietal lobule (IPL).

|  | HCF/low<br>ADNC | HCF/high ADNC | Sporadic<br>ADD | Autosomal Dominant<br>ADD |
| --- | --- | --- | --- | --- |
| N | 8 | 12 | 18 | 23 |
| Age (*mean ± sd*) | 89 ± 4 | 89 ± 6 | 80 ± 13 | 50 ± 10 |
| Sex (*M:F*) | 4:4 | 6:6 | 11:7 | 15:8 |
| Post mortem interval<br>hr (*mean ± sd*) | 3.9 ± 0.9 | 4.9 ± 1.5 | 4.5 ± 1.2 | 15.7 ± 9.3 |
| APOE ε4 alleles (*n (%)*) | 2 (12.5%) | 6 (20.8%) | 10 (27.8%) | 6 (18.8%)* |
| ε3:ε4 (*n*) | 2 | 4 | 6 | 4 |
| ε4:ε4 (*n*) | 0 | 1 | 2 | 1 |
| *missing* | 0 | 0 | 0 | 7 |
| Mutations (*n x gene*) | NA | NA | NA | 17 x *PSEN1*, 5 x *PSEN2*,<br>1 X *APP* |
| Braak Stage |  |  |  |  |
| B1 (*n*) | 3 | 0 | 0 | 0 |
| B2 (*n*) | 5 | 6 | 0 | 0 |
| B3 (*n*) | 0 | 6 | 18 | 23 |
| CERAD Score |  |  |  |  |
| C0 (*n*) | 8 | 0 | 0 | 0 |
| C2 (*n*) | 0 | 6 | 4 | 1 |
| C3 (*n*) | 0 | 6 | 14 | 22 |

\*For these; if there were missing genotypes the percentage is of the reported genotypes

**Table 4.** Brain donor characteristics for the caudate nucleus.

|  | HCF/low ADNC | HCF/high ADNC | Sporadic ADD | Autosomal Dominant ADD |
| --- | --- | --- | --- | --- |
| N | 10 | 10 | 18 | 20 |
| Age (*mean ± sd*) | 85 ± 6 | 90 ± 5 | 81 ± 11 | 51 ± 11 |
| Sex (*M:F*) | 6:4 | 5:5 | 10:8 | 13:7 |
| Post mortem interval hr (*mean ± sd*) | 3.9 ± 0.8 | 5.2 ± 1.4 | 4.4 ± 1.3 | 15.2 ± 9.8 |
| *APOE* ε4 alleles (*n (%)*) | 2 (10.0%) | 5 (25.0%) | 9 (25.0%) | 5 (17.9%)* |
| ε3:ε4 (*n*) | 2 | 3 | 5 | 3 |
| ε4:ε4 (*n*) | 0 | 1 | 2 | 1 |
| *missing* | 0 | 0 | 0 | 6 |
| Mutations (*n x gene*) | NA | NA | NA | 17 x *PSEN1*, 6 x *PSEN2*, 1 X *APP* |
| Braak Stage |  |  |  |  |
| B1 (*n*) | 6 | 0 | 0 | 0 |
| B2 (*n*) | 4 | 5 | 0 | 0 |
| B3 (*n*) | 0 | 5 | 18 | 20 |
| CERAD Score |  |  |  |  |
| C0 (*n*) | 10 | 0 | 0 | 0 |
| C2 (*n*) | 0 | 5 | 4 | 1 |
| C3 (*n*) | 0 | 5 | 14 | 19 |

\*For these; if there were missing genotypes the percentage is of the reported genotypes

### Ethics oversight

All study cohort participants were collected and provided informed consent under protocols approved by the Institutional Review Board (IRB) at University of Washington, Kaiser Permanente Washington, and Stanford University. The UW and Stanford Human Subjects Division deems the use of pre-existing de-identified samples exempt from full IRB review and, thus, treated this project as non-human subjects research.

### Sample metadata, batch design and references

Each human brain region was divided into batches of 14 individual samples and 2 pooled references for a total of 16. The first batch of each region was also used to create a region-specific reference pool to be used as a “common reference” and/or single point calibrant, which was homogenized, aliquoted, frozen, and used to compare between batches within a brain region. Human cerebellum and occipital lobe tissue was homogenized, pooled, aliquoted and frozen to be used as a “batch reference” for comparison between batches and other brain regions. Batch design was randomly balanced based on group ratios. For example, batches from the SMTG brain region contained 5 “Sporadic ADD”, 4 “Autosomal Dominant ADD”, 2 “HCF/low ADNC”, and 3 “HCF/high ADNC” samples. Metadata for the samples from the SMTG, Hippocampus, IPL and Caudate brain regions is provided in Supplementary Tables 1-4 available on Panorama Public in the “Supplementary Data” subfolder^15,16^. For each region the metadata includes; sample batch, age, sex, post-mortem interval, APOE genotype, cognitive status, study of origin, and consensus Braak stage and CERAD score.

### Sample homogenization and protein digestion

Two 25 μm frozen sections of brain tissue were resuspended in 120 μl of lysis buffer of 5% SDS, 50mM triethylammonium bicarbonate (TEAB), 2mM MgCl2, 1X HALT phosphatase and protease inhibitors, vortexed and briefly sonicated at setting 3 for 10 s with a Fisher sonic dismembrator model 100. A microtube was loaded with 30 μl of lysate and capped with a micropestle for homogenization with a Barocycler 2320EXT (Pressure Biosciences Inc.) for a total of 20 minutes at 35°C with 30 cycles of 20 seconds at 45,000 psi followed by 10 seconds at atmospheric pressure. Protein concentration was measured with a BCA assay. Homogenate of 50 μg was added to a process control of 800 ng of yeast enolase protein (Sigma) which was then reduced with 20 mM DTT and alkylated with 40 mM IAA. Lysates were then prepared for S-trap column (Protifi) binding by the addition of 1.2% phosphoric acid and 350 μl of binding buffer (90% Methanol, 100 mM TEAB). The acidified lysate was bound to column incrementally, followed by 3 wash steps with binding buffer to remove SDS and 3 wash steps with 50:50 methanol:chloroform to remove lipids and a final wash step with binding buffer. Trypsin (1:10) in 50mM TEAB was then added to the S-trap column for digestion at 47°C for one hour. Hydrophilic peptides were then eluted with 50 mM TEAB and hydrophobic peptides were eluted with a solution of 50% acetonitrile in 0.2% formic acid. Elutions were pooled, speed vacuumed and resuspended in 0.1% formic acid.

Injection of samples are one ug of total protein (16 ng of enolase process control) and 150 fmol of a heavy labeled Peptide Retention Time Calibrant (PRTC) mixture (Pierce). The PRTC is used as a peptide process control. Library pools are an equivalent amount of every sample (including references) in the batch. For example, a batch library pool consists of the 14 samples from the batch and two references. System suitability (QC) injections are 150 fmol of PRTC and BSA.

### Liquid chromatography and mass spectrometry

One µg of each sample with 150 femtomole of PRTC was loaded onto a 30 cm fused silica picofrit (New Objective) 75 µm column and 3.5 cm 150 µm fused silica Kasil1 (PQ Corporation) frit trap loaded with 3 µm Reprosil-Pur C18 (Dr. Maisch) reverse-phase resin analyzed with a Thermo Easy-nLC 1200. The PRTC mixture is used to assess system suitability before and during analysis. Four of these system suitability runs are analyzed prior to any sample analysis and then after every six sample runs another system suitability run is analyzed. Buffer A was 0.1% formic acid in water and buffer B was 0.1% formic acid in 80% acetonitrile. The 40-minute system suitability gradient consists of a 0 to 16% B in 5 minutes, 16 to 35% B in 20 minutes, 35 to 75% B in 1 minute, 75 to 100% B in 5 minutes, followed by a wash of 9 minutes and a 30-minute column equilibration. The 110-minute sample LC gradient consists of a 2 to 7% for 1 minutes, 7 to 14% B in 35 minutes, 14 to 40% B in 55 minutes, 40 to 60% B in 5 minutes, 60 to 98% B in 5 minutes, followed by a 9 minute wash and a 30-minute column equilibration. Peptides were eluted from the column with a 50°C heated source (CorSolutions) and electrosprayed into a Thermo Orbitrap Fusion Lumos Mass Spectrometer with the application of a distal 3 kV spray voltage. For the system suitability analysis, a cycle of one 120,000 resolution full-scan mass spectrum (350-2000 *m*/*z*) followed by a data-independent MS/MS spectra on the loop count of 76 data-independent MS/MS spectra using an inclusion list at 15,000 resolution, AGC target of 4e5, 20 millisecond (ms) maximum injection time, 33% normalized collision energy with a 8 *m*/*z* isolation window. For the sample digest, first a chromatogram library of 6 independent injections is analyzed from a pool of all samples within a batch. For each injection a cycle of one 120,000 resolution full-scan mass spectrum with a mass range of 100 *m*/*z* (400-500 *m*/*z*, 500-600 *m*/*z*, 600-700 *m*/*z*, 700-800 *m*/*z*, 800-900 *m*/*z*, 900-1000 *m*/*z*) followed by a data-independent MS/MS spectra on the loop count of 26 at 30,000 resolution, AGC target of 4e5, 60 ms maximum injection time, 33% normalized collision energy with a 4 *m*/*z* overlapping isolation window. The chromatogram library data is used to quantify proteins from individual sample runs. These individual runs consist of a cycle of one 120,000 resolution full-scan mass spectrum with a mass range of 350-2000 *m*/*z*, AGC target of 4e5, 100 ms maximum injection time followed by a data-independent MS/MS spectra on the loop count of 76 at 15,000 resolution, AGC target of 4e5, 20 ms maximum injection time, 33% normalized collision energy with an overlapping 8 *m*/*z* isolation window. Application of the mass spectrometer and LC solvent gradients are controlled by the ThermoFisher Xcalibur (version 3.1.2412.24) data system. Mass spectrometry run order for all samples is provided in Supplementary Tables 5-8 available on Panorama Public.

### Peptide detection and quantitative signal processing

Thermo RAW files were converted to mzML format using Proteowizard (version 3.0.20064) using vendor peak picking and demultiplexing with the settings of “overlap_only” and Mass Error = 10.0 ppm^5^. On column chromatogram libraries were created using the data from the six gas phase fractionated “narrow window” DIA runs of the pooled reference as described previously^17^. These narrow windows were analyzed using EncyclopeDIA (version 1.4.10) with the default settings (10 ppm tolerances, trypsin digestion, HCD b- and y-ions) of a Prosit predicted spectra library based the Uniprot human canonical FASTA (January 2021). The results from this analysis from each brain region were saved as a “Chromatogram Library’’ in EncyclopeDIA’s eLib format where the predicted intensities and iRT of the Prosit library were replaced with the empirically measured intensities and RT from the gas phase fractionated LC-MS/MS data. The “wide window” DIA runs were analyzed using EncyclopeDIA (version 1.4.10) requiring a minimum of 3 quantitative ions and filtering peptides with q-value ≤ 0.01 using Percolator 3.01. After analyzing each file individually, EncyclopeDIA was used to generate a “Quant Report’’ which stores all the detected peptides, integration boundaries, quantitative transitions, and statistical metrics from all runs in an eLib format. The Quant Report eLib library is imported into Skyline (daily version 22.2.1.278) with the human uniprot FASTA as the background proteome to map peptides to proteins, perform peak integration, manual evaluation, and report generation. A csv file of peptide level total area fragments (TAFs) for each replicate was exported from Skyline using the custom reporting capabilities of the document grid^18^.

### Quantitative data post-processing, normalization, and batch correction

Despite precautions taken to ensure equivalent sample preparation, handling and acquisition, additional post-processing was performed to normalize, and batch adjust the quantitative data to remove residual technical noise. Modeling the proportional changes of peptide/protein group intensities, log_2_ transformation is applied followed by a Median Deviation (MD) normalization to the peptide total area fragment values (level 2 data) across instrument runs within a brain region (Equation 1) under the assumption that median total area fragment values should be equal sans batch effect from known and unknown sources of variability.

MD Normalization, by calculating the deviation from the median of the sample total area fragment, should neither remove scale information nor de-weigh outlier signals that may be of biological relevance^19^. Here, the MD normalized peptide *F* of each sample is given by the following. The peak areas (*A*_*i*_) for each peptide *i* are first log_2_ transformed and then normalized by equalizing the median peak areas across all samples using the equation:

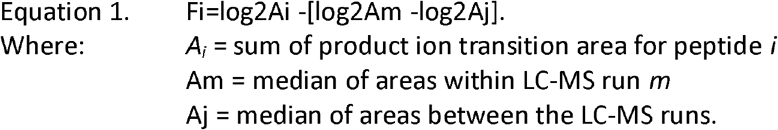

The effectiveness and validity of the normalization approach is then assessed by evaluating the comparability of the peptide abundance distribution across samples (Figure 2B), and by the reproducibility of those peptide abundances across replicate samples (Figure 3A). Peptide abundances are then adjusted for batch effect by fitting a linear model and “regressing” out the factors with known unwanted sources of variation to return a matrix of residuals. The detection of the presence of batch effect pre- and post-adjustment is assessed by exploring the data variance structure through Principal Variance Component Analysis (PVCA) (https://bioconductor.org/packages/release/bioc/html/pvca.html) (Supplementary Figure 4 available on Panorama Public) and Principal component Analysis (PCA) using projections onto the first three principal components. The normalization and batch adjusted peptide abundances are available as the level 3A data file. Using DIA, all observable peptides in one sample will be sampled in all of the other biological replicates^12,13,20^. Due to the comprehensive sampling nature of DIA we can extract information for the same transitions across all samples in an experiment. The resulting zeros in our peptide abundance data therefore represent signals below our limit of detection and are not treated as missing data. After protein group inference, protein abundances are batch corrected using the same method as the peptide data.

### Protein grouping and inference

The processing and ‘roll-up’ of DIA data borrows from the established strategies adopted in the DDA field in which the quantification of peptides and their corresponding protein groups is inferred through the modification of IDPicker algorithm^21^. In summary, to quantify the peptide/protein groups, a bipartite graph of peptide-protein interactions is constructed to generate groupings through the parsimony reduction of the graph as it is implemented in MSDaPl. Then, the peptide abundances at the nodes are summed to estimate the abundance of the peptide groups and proteins that match the same set of peptide groups are merged into a single node in the graph, forming an indistinguishable protein group^22^.

## Data Records

The Skyline documents, raw files for quality control and DIA data are available at Panorama Public. ProteomeXchange ID: PXD034525. DOI: https://doi.org/10.6069/wefm-vv52. Access URL: https://panoramaweb.org/ADBrainCleanDiagDIA.url^16^.

DIA data is available in 5 different categories based on the level of post-processing (Figure 1e) for each brain region. Level 0 represents the raw data in two different formats - the raw format is directly from the Thermo mass spectrometer and the mzML format is the demultiplexed version of the raw data (Proteowizard version 3.0.20064). Level 1 describes the zipped Skyline document grouped by batch. Level 2 is a csv file grouped by batch of the Skyline output with the integrated peak area for each peptide (row) in each replicate (column). Level 3A is a csv file of the normalized peptide abundance across all batches. Level 3B is a csv file of the normalized protein abundance across all batches.

**Figure 1:**
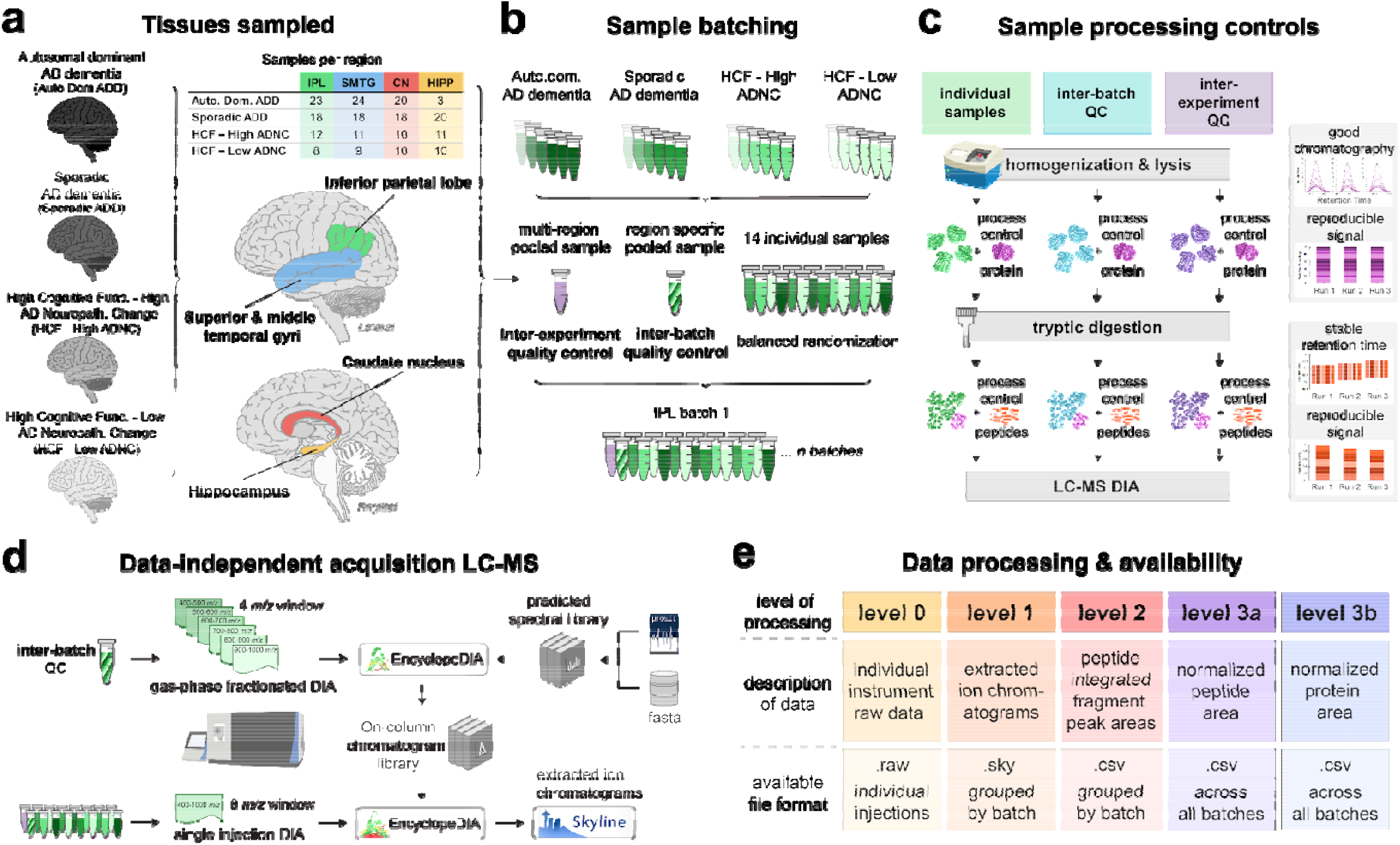
Experimental scheme for the collection of the proteomics data using data independent acquisition-mass spectrometry. A) Brain tissue sections from 4 regions were analyzed for all groups. B) Samples were prepared and analyzed in batches of 16, with 14 individual samples per batch selected by balanced randomization. Each batch contained an inter-experiment quality control sample generated from pooling portions of several individual samples from across all 4 brain regions sampled. Each batch also contained an inter-batch quality control sample generated from pooling portions of individual samples within that brain region. C) In addition to quality control samples, both protein and peptide sample processing controls were included in all samples to track system suitability. D) For each batch an on-column data-independent acquisition chromatogram library is generated from overlapping, narrow window gas-phase fractionation of an inter-batch QC. Individual samples are acquired by a single injection wider window data-independent acquisition method. Peptide detection and scoring is performed using EncyclopeDIA and extracted ion chromatograms integrated with Skyline. E) The proteomics data is publicly available on the Panorama web server in 5 different states, each corresponding to the level of post-processing.

Quality control Skyline documents, peptide QC plots and instrument raw files for system suitability runs are provided by brain region. The Skyline documents and peptide QC plots for enolase and PRTC process controls are provided by brain region. The instrument raw files for process controls are the same as DIA sample raw files by brain region. An interactive dashboard is available for the SMTG and Hippocampus data.

## Technical Validation

### Balanced and controlled experiment design

We have designed our experiment to perform quantitative, peptide-centric proteomics using brain tissue from four different brain regions selected specifically because they represent distinct anatomical regions with varying pathological involvement by AD (Figure 1). The experimental design was intended to compare individual samples from the four different categorical disease groups within each brain region. Samples were prepared in batches of 16 samples which consisted of 14 brain tissue samples and two external control samples. The batch size was determined by the number of samples, 16, that could be prepared within a Barocycler (Pressure Biosciences, Inc.). For each batch, the samples were randomized in a balanced block design (Supplementary Table 5 available on Panorama Public).

Within each batch we included both internal and external controls. Internal controls were added to each sample to provide a QC check of the sample preparation and LC-MS data collection process. These “Process Controls” consisted of the addition of yeast enolase protein after lysis and prior to digestion and the Pierce Retention Time Calibration (PRTC; 15 synthetic stable isotope labeled peptides) peptide mixture following digestion. The “Protein Internal Control” was used to assess the protein digest and peptide recovery and the “Peptide Internal Control” was used to distinguish between sample preparation and measurement issues post-digestion.

The two external controls were different brain lysates that were prepared, measured, and analyzed with the rest of the samples in the batch. One of the controls was a brain region specific pool used to assess between batch quality control. This inter-batch quality control is composed of a randomized balanced pooled sample set for each respective brain region. For example, the inter-batch quality control “TRPR” is composed of 3 HCF/high ADNC samples, 3 HCF/low ADNC samples, 3 AutoDom ADD samples and 5 Sporadic ADD samples from the SMTG. The same inter-batch quality control is run in every batch of the experiment from the SMTG and was used to assess data quality post-normalization. The second external control was an inter-brain region quality control (“HAD” samples) and composed of a homogenate of a mix of cerebellum and occipital lobe tissue which we had ample material available to use throughout all our brain tissue experiments. The cerebellum and occipital lobe external control is distinct from the rest of the brain tissue regions in the experiment, but this should not hinder the interpretation of the experimental results, as this control is only monitoring the reproducibility of our entire system. The same pool of inter-brain external control was prepared and run in every batch across all brain regions for the entire experiment.

We can determine when our sample preparation and system is not functioning as expected with a combination of system suitability checks, inter-batch quality controls, inter-experiment quality controls and process controls (Figure 1b). Our system suitability consists of a mixture of a BSA tryptic digest and PRTC prior to sample analysis and throughout sample collection at a frequency of once every six to eight samples.

### Run level and experiment level peptide and protein detections

For each sample in each brain region several tryptic peptides can be detected at a 1% FDR cut-off. IPL samples ranged from 37840-73168, SMTG ranged from 51582-69590, hippocampus from 32995-59853, and caudate nucleus ranged from 31426-58105 (Table 5). To integrate data across all individual samples within each brain region we control with an experiment level error rate. This leads to the same peptides quantified in all samples within a brain region; 48271 in IPL, 40346 in SMTG, 31863 in hippocampus, and 26135 in caudate nucleus. These peptides map to 6497 quantified proteins in IPL, 5851 in SMTG, 5117 in hippocampus, and 4636 in caudate nucleus (Table 5). The distribution of peptide abundances is aligned with median normalization, as demonstrated with the SMTG data (Figure 2B) as well as all brain regions (Supplementary Figures 1 and 2 available on Panorama Public).

**Table 5.**
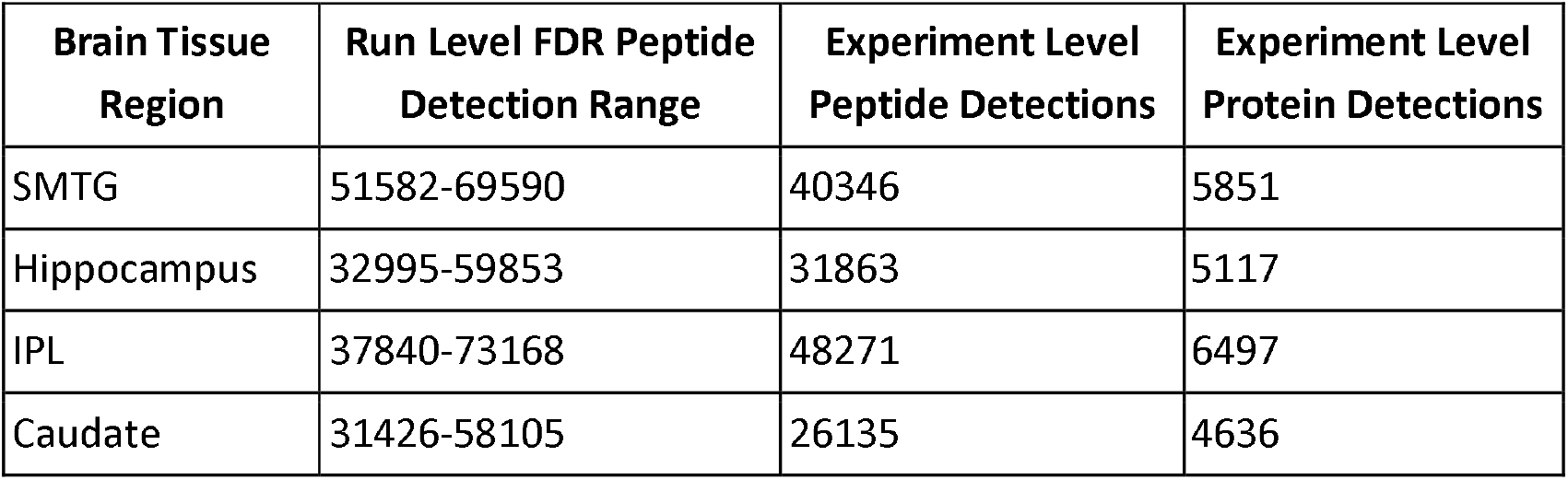
Run Level and Experiment Level Peptide and Protein Detections.

**Figure 2:**
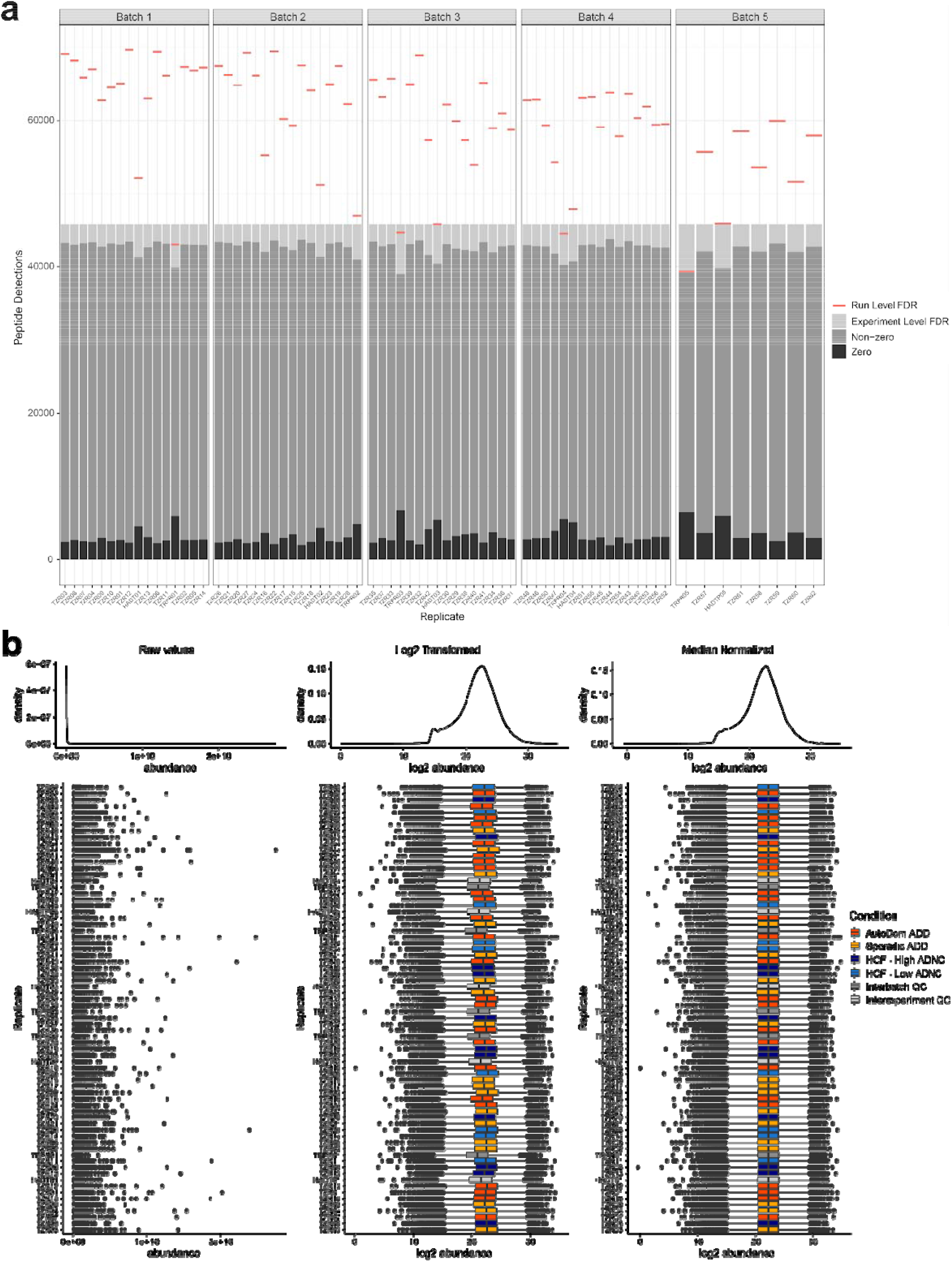
Peptides detected in individual runs and across the entire dataset for SMTG. A) The number of peptides detected for each sample in the SMTG brain region experiment, with those detected in the sample at a 1% FDR indicated by the red line, and those detected across all samples at an 1% experiment level FDR indicated in gray. Although zeros exist in this remaining data, these represent a lack of signal above background. B) Kernel density plots (above) and box plots (below) for each individual sample show the distribution of peptide abundances before and after log2 transformation and then median normalization. Sample groups are highlighted with different colors in the log2 transformation and median normalization box plots. Figure is generated with level 2 data.

### Inter-batch precision and reproducibility

The inclusion of inter-batch quality control samples allows us to assess the impact of normalization and batch correction on peptide and protein quantitative reproducibility. For example, the SMTG experiment was split into 5 batches for processing and acquisition. Using the inter-batch control replicate samples, the coefficient of variation can be calculated for all peptides quantified in the SMTG. The distribution of peptide coefficient of variation improves with normalization and batch correction, with the mean decreasing about 8.2%. Likewise, the protein coefficient of variation also improves following batch correction, with the mean decreasing by about 1.25% (Figure 3). Peptide and protein quantities are highly correlated across inter-batch replicates. Inter-batch control replicate samples in SMTG have peptide Pearson correlation coefficients ranging from 0.867 to 0.950, and protein correlations ranging from 0.894 to 0.960 (Figure 4).

**Figure 3:**
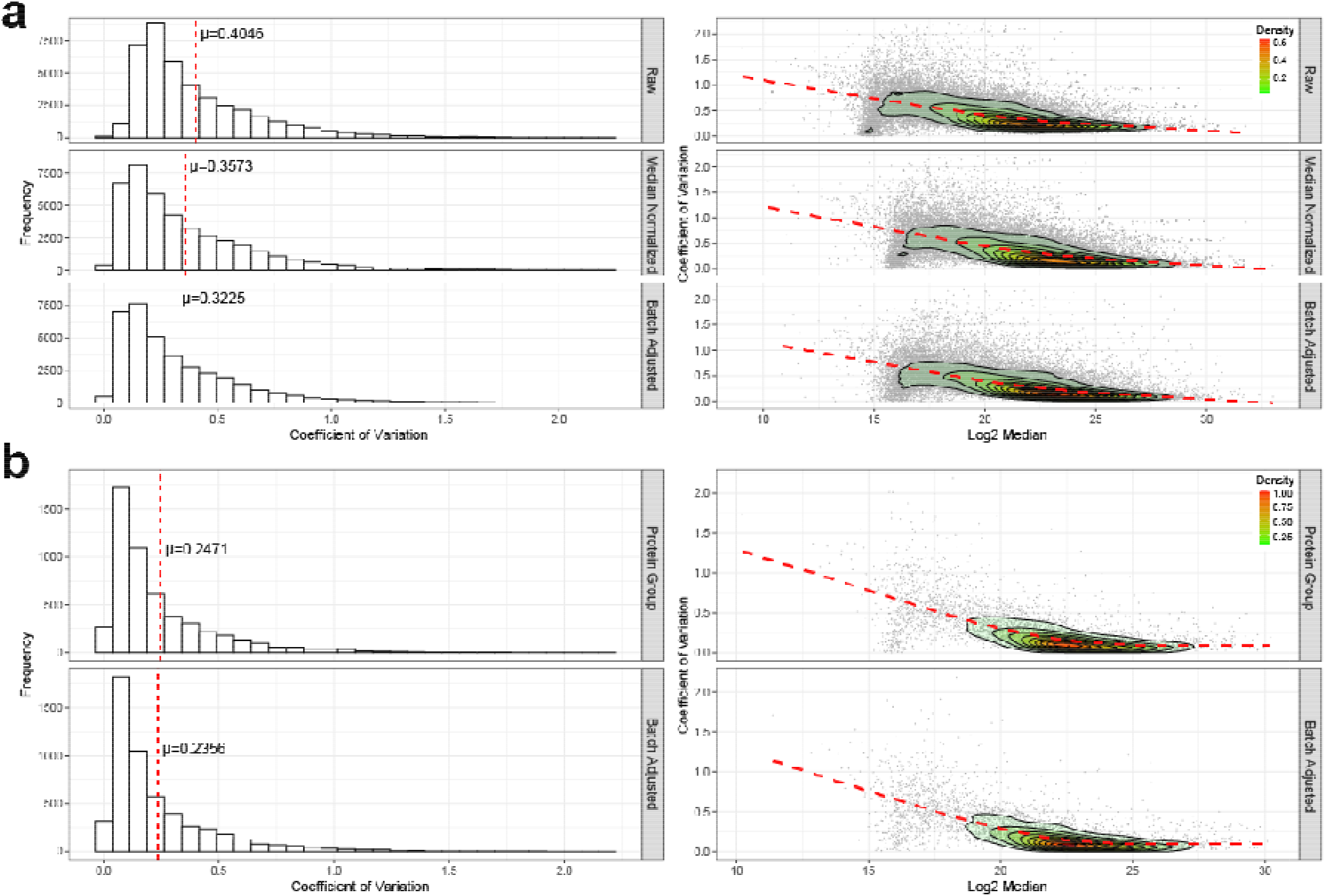
Effect of normalization and batch correction on the inter-batch variance for SMTG. A) The effect of median normalization and batch correction on the inter-batch peptide coefficient of variance. Mean coefficient of variation (μ) is indicated by the red line. Figure 3a is generated from level 2 to 3a data. B) Effect of batch correction on the inter-batch protein coefficient of variance. The relationship between coefficient of variation and the log2 median abundance is visualized with a loess fit of a contoured density plot (red line). Figure 3b is generated from level 2 to 3b data.

**Figure 4:**
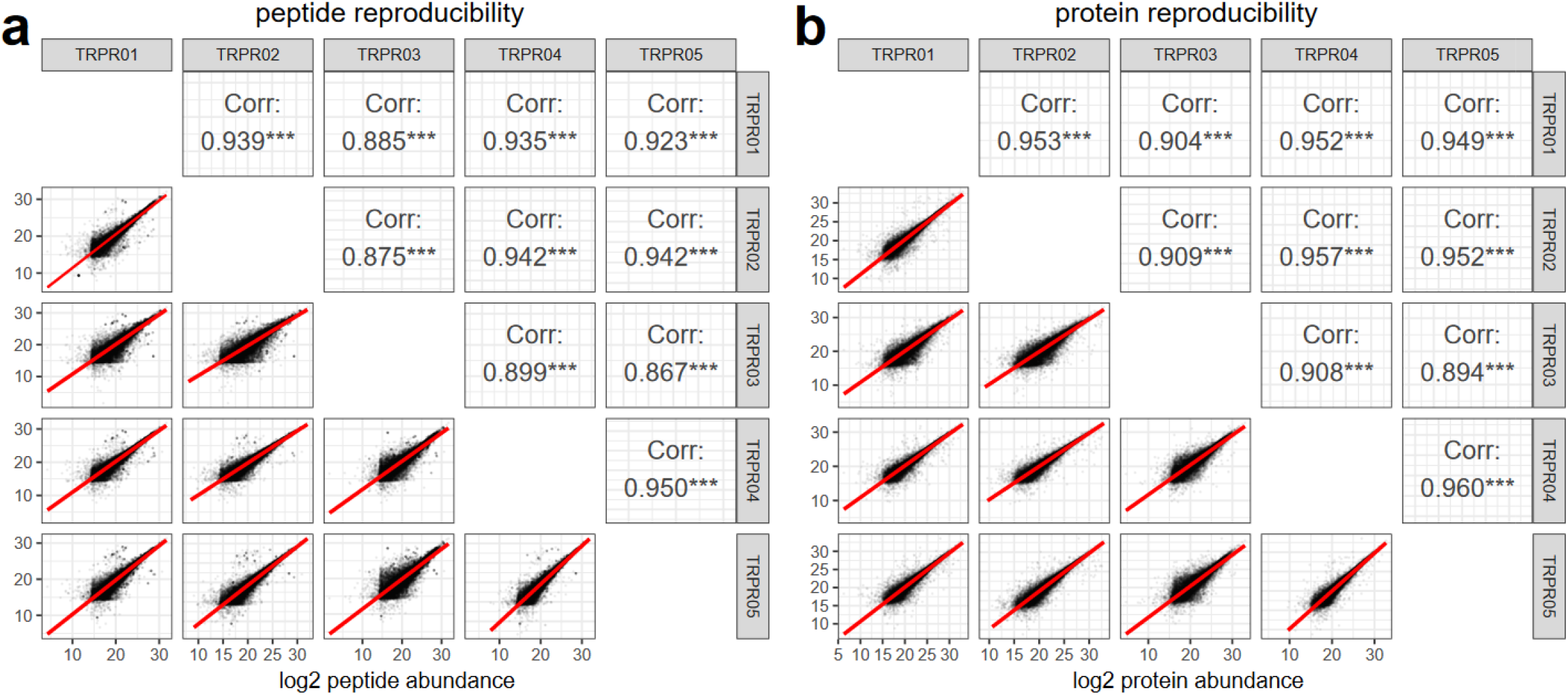
Correlation of the quantitative results from five different sample preparation and analysis batches following normalization and batch correction for SMTG. A) The log2 peptide abundances measured in the inter-batch quality control samples across each of the 5 SMTG experimental batches, with the Pearson correlation coefficient. Figure 4a is generated from level 3a data. B) The log2 protein group abundances from the inter-batch quality control samples in the 5 SMTG batches samples, and their Pearson correlation coefficient. Significance is indicated with *** being p < 0.001. Figure 4b is generated from level 3b data.

**Figure 5:**
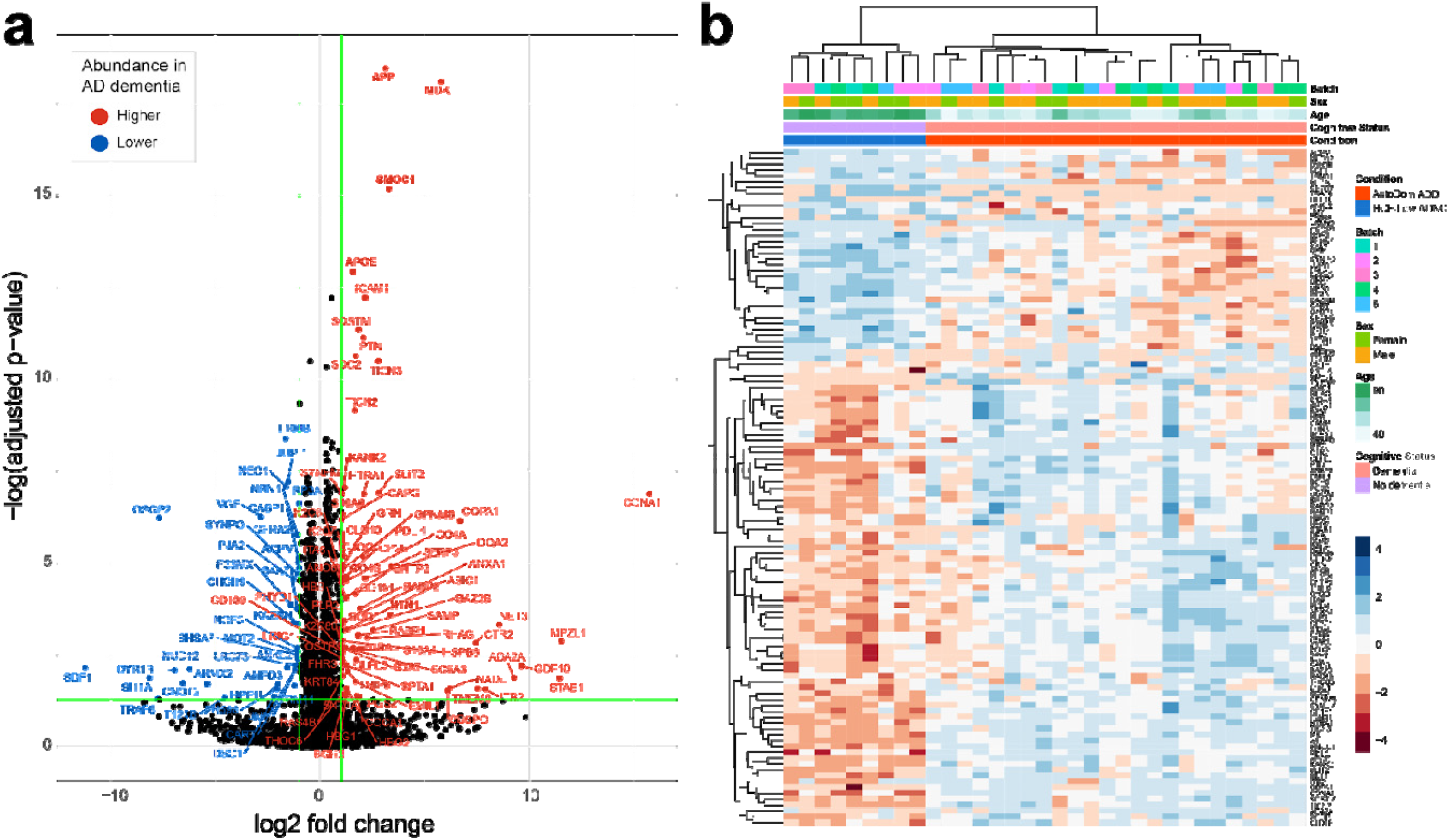
Summary data of quantitative changes observed for SMTG. A) Volcano plot showing proteins that are statistically different between the autosomal dominant Alzheimer’s disease (AD) dementia (ADD) and the high cognitive function (HCF)/low AD neuropathologic change (ADNC) SMTG using cut-off of log2 fold-change > ± 0.5 and FDR adjusted p-value < 0.05. B) Heatmap showing complete linkage hierarchical clustering of 33 SMTG samples using the 115 proteins with significant differences between ADD and the HCF-low ADNC samples. Figure 5 is generated from level 3b data.

### Expected biological differences

Of the peptides quantified, we detect several proteins known to be involved in AD. In the SMTG data we quantify peptides mapping to 250 proteins found in AD-related pathways (Supplemental Figure 5 available on Panorama Public). Preliminary assessment of peptide and protein data show we can distinguish between sample groups (Supplemental Figure 3 available on Panorama Public). Differential abundance analysis between HCF/low ADNC and autosomal dominant ADD in SMTG captures known biology. Protein groups found to be significantly different between the groups include previously documented increases in Amyloid precursor protein (APP/A4), Apolipoprotein E (APOE), SPARC-related modular calcium-binding protein 1 (SMOC1), midkine (MK/MDK) and netrin-1 (NET1/NTN1)^23^. We detect and quantify two peptides mapping to the n- and c-terminal sides of the alpha-secretase cleavage site in amyloid-β sequence. Both peptides have an expected difference in abundance across the sample groups in all four brain regions, with autosomal dominant ADD having the highest distribution, followed by sporadic ADD, HCF/high ADNC, and HCF/low ADNC. Peptides mapping to microtubule-associated protein tau (MAPT) also have some expected differences between the experimental groups. In the SMTG brain region the seven quantified peptides spanning the microtubule binding region of MAPT (residues 243-368) are increased in autosomal dominant ADD, followed by sporadic AD, compared to both HCF groups. This trend has been observed previously and leads to the aggregated protein abundance to be differential in the same manner as the microtubule binding region peptides^24^.

## Usage Notes

We provide quantitative data both for tryptic peptides directly and for protein groups derived from peptide quantities. Based on extensive existing knowledge of AD, we know that modified forms of the APP or amyloid-β and Tau proteins are important in disease progression. Historically mass spectrometry proteomics data has aggregated multiple tryptic peptides per protein coding sequence to arrive at a singular protein level value. This would result in a loss of important information regarding the status of the individual peptides^7^. By collecting these samples by DIA we can reproducibly quantify individual tryptic peptides across our entire sample set. With this peptide data we observe differentially abundant peptides within a protein coding gene. If aggregated to a singular value this important signal would be lost.

### Use Case 1. Sporadic and autosomal dominant Alzheimer disease dementia differential peptide and protein abundance

This mass spectrometry data has typically been reported as relative protein group abundance measures, enabling differential abundance testing of protein groups between disease states (Figure 6). In SMTG there are proteins with statistically different abundance between healthy controls and AD. This type of analysis can be extended to all brain regions. A large portion of research investigating the molecular and pathological basis of AD has been generated using model systems informed by genetic causes. Studying individuals with sporadic ADD can be challenging due to the presence of comorbidities. Here we present data generated from both autosomal dominant ADD with minimal comorbidities and sporadic ADD with minimal comorbidities. These data can be used to better understand biological differences or similarities in these two types of ADD.

**Figure 6:**
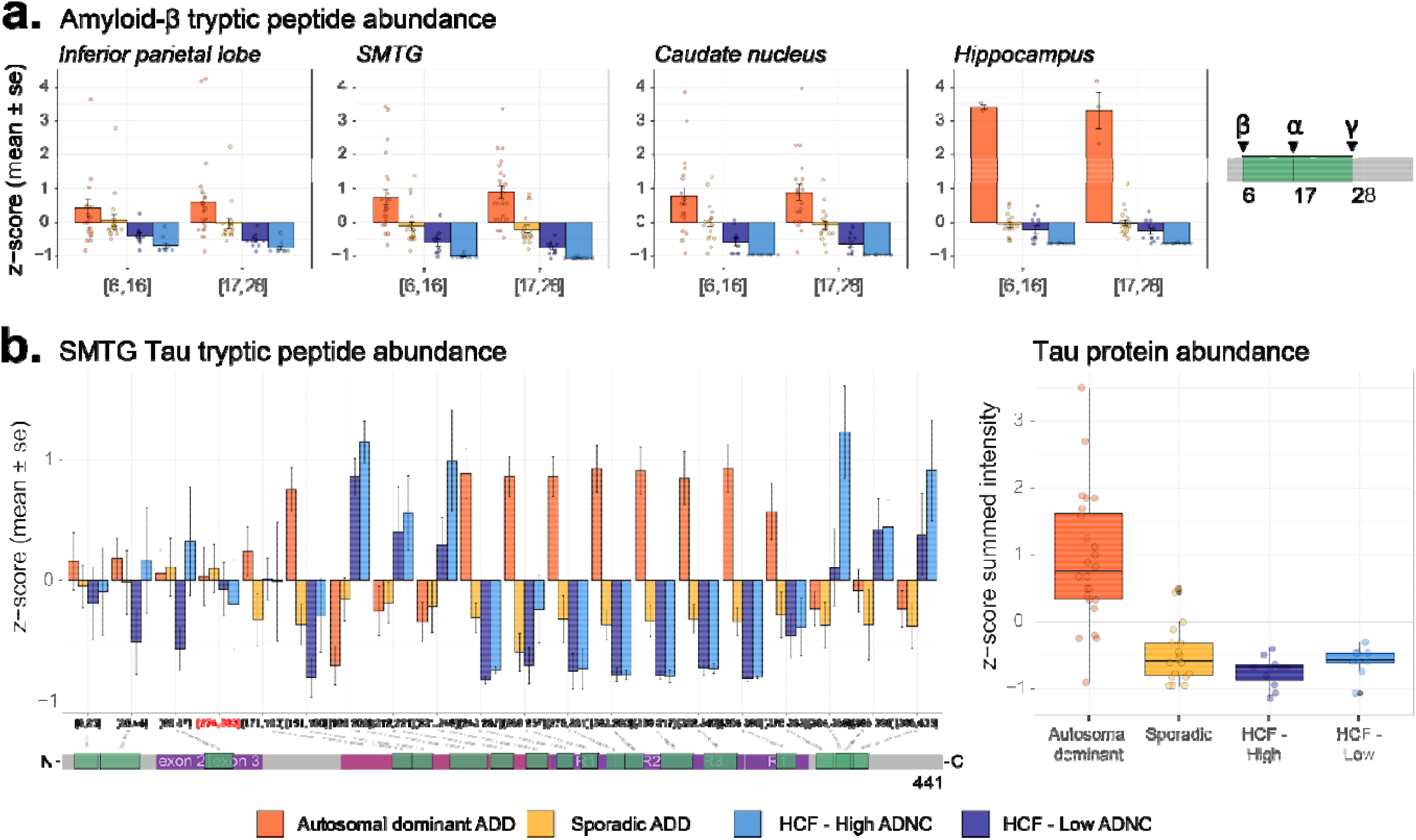
Amyloid-β and tau peptide abundances recapitulate known molecular changes for the SMTG. A) Two tryptic peptides mapping to amyloid-β (amino acid residues 6-16: HDSGYEVHHQK, 17-28: LVFFAEDVGSNK) are z-scored across all individuals and plotted as the mean and standard error for each sample group. B) Abundances for tryptic peptides mapping to tau are z-scored across all individuals and plotted as the mean and standard error for each sample group. While there are some common signatures within protein domains, these signatures are lost if aggregated to a protein level. Tryptic peptides are labeled based on their first and last amino acid residues. Figure 6 is generated from level 3a data.

### Use Case 2. Differential peptide and protein abundance with neuropathologic markers and dementia

The HCF/high ADNC samples have classic histopathologic features of AD - amyloid-β plaques and neurofibrillary tangles - at levels overlapping with individuals who have Sporadic ADD. This enables analyses to measure the differences in protein pathology between individuals who have developed dementia and those who have not developed dementia despite similar levels of high amyloid-β plaques and Tau tangles.

### Use Case 3. Mass spectrometry data reuse and reanalysis

Beyond the analyses presented here, the storage of this data is done in a way that facilitates use of this data for informing subsequent mass spectrometry assay development. All data from this project is available on the Panorama server-based data repository application for targeted mass spectrometry assays^15,16^. Thus, all extracted ion chromatogram information from every sample across all brain regions is available in an interactive Skyline document format, and readily available for download and reuse. Information about fragment ions and chromatography is important for the development of targeted assays^25,26^, making this data valuable to the wider community.

Using DIA mass spectrometry methods allows for the data to be easily reanalyzed. This could be used to search for post-translational modifications or sequence variants not searched for in our current analysis. Additionally, this data can be reanalyzed to look at other unique features, such as isomerized peptides. The feasibility of this was recently shown by Hubbard et al. through reanalysis of a subset of this dataset^27^. Peptide-centric reanalysis is possible due to the comprehensive sampling by data-independent acquisition of a preset range across all samples.

## Code Availability

The MSConvert installer and documentation is available from https://proteowizard.sourceforge.io/. EncyclopeDIA is available at https://bitbucket.org/searleb/encyclopedia/. The Skyline-daily installer and documentation are available from https://skyline.ms/skyline.url. The source code for both the MSConvert and Skyline projects are available as part of the Proteowizard project https://github.com/ProteoWizard/pwiz

All code used for the analysis of the data matrix including data QC, normalization, pre-processing, visualization, and figure generation is available online at https://github.com/uw-maccosslab/ADBrainCleanDiagDIA.

## Acknowledgements

Support for this research was provided by the National Institute of Aging (RF1 AG053959, U19 AG065156, F31 AG069420, R01 AG075802). The collection of tissue samples were supported by the UW Alzheimer’s Disease Research Center (P30 AG066509), the Adult Changes in Thought study U19 AG066567, the Nancy and Buster Alvord Endowment (C.D.K.), the Massachusetts Alzheimer Disease Research Center (P30 AG062421), The Dominantly Inherited Alzheimer Network (U19 AG032438, UF1AG032438), the German Center for Neurodegenerative Diseases (DZNE), Raul Carrea Institute for Neurological Research (FLENI), the Research and Development Grants for Dementia from Japan Agency for Medical Research and Development, AMED, and the Korea Health Technology R&D Project through the Korea Health Industry Development Institute (KHIDI). This manuscript has been reviewed by DIAN Study investigators for scientific content and consistency of data interpretation with previous DIAN Study publications. We acknowledge the altruism of the participants and their families and contributions of the research and support staff at each of the participating sites for their contributions to this study.

## Author contributions

G.E.M., J. P., and D.P. share equal first authorship. G.E.M., M.J.M., and T.J.M. designed the experiment. G.E.M. prepared the brain tissue, ran the mass spectrometer, performed the bioinformatic signal processing, and organized the data repository. J.P. performed data quality control, normalization, batch correction and statistics. J.P. and D.P. produced the figures and interpreted the results. B.C.S. designed mass spectrometry methods and software. C.D.K., E.B.L., R.B., R.J.P., J.P.C., M.R.F., C.A.M., B.G., K.L.N. and M.P.F. provided specimens. G.E.M., J.P., D.P., T.J.M. and M.J.M. wrote the paper. All authors reviewed the manuscript.

