## Supplementary Tables for "A peptide-centric quantitative proteomics dataset for the phenotypic assessment of Alzheimer’s disease"

**Supplementary Table 1. Summary metadata table for the superior and middle temporal gyri (SMTG).**

| **Batch** | **Sample Label** | **Condition** | **Age** | **Sex** | **Mutation Status** | **PMI (hrs)** | ***APOE* Alleles** | **Cognitive Status** | **Study Name** | **Braak Stage** | **CERAD Score** |
| --- | --- | --- | --- | --- | --- | --- | --- | --- | --- | --- | --- |
| 1 | TZR01 | Autosomal Dominant ADD | 52 | Female | PSEN1 | NA | 3_4 | Dementia | UW ADRC | 3 | 3 |
| 1 | TZR02 | Sporadic ADD | 90 | Female | NA | 3.85 | 3_3 | Dementia | ACT | 3 | 3 |
| 1 | TZR03 | Sporadic ADD | 61 | Male | NA | 3 | 4_4 | Dementia | UW ADRC | 3 | 3 |
| 1 | TZR04 | Sporadic ADD | 63 | Male | NA | 3.33 | 3_3 | Dementia | UW ADRC | 3 | 3 |
| 1 | TZR05 | Sporadic ADD | 77 | Male | NA | 5.92 | 3_3 | Dementia | ACT | 3 | 3 |
| 1 | TZR06 | HCF/High ADNC | 93 | Male | NA | 6.83 | 3_3 | No dementia | ACT | 3 | 2 |
| 1 | TZR07 | Autosomal Dominant ADD | 39 | Female | PSEN1 | 25 | NA | Dementia | DIAN | 3 | 3 |
| 1 | TZR08 | HCF/High ADNC | 89 | Female | NA | 2.5 | 3_4 | No dementia | ACT | 3 | 3 |
| 1 | TZR09 | Sporadic ADD | 88 | Male | NA | 4.38 | 3_3 | Dementia | UW ADRC | 3 | 3 |
| 1 | TZR10 | Autosomal Dominant ADD | 44 | Female | PSEN1 | 17.5 | 3_3 | Dementia | UW ADRC | 3 | 3 |
| 1 | TZR11 | HCF/Low ADNC | 91 | Male | NA | 5 | 3_3 | No dementia | ACT | 2 | 0 |
| 1 | TZR12 | Autosomal Dominant ADD | 78 | Female | PSEN2 | 7 | 2_3 | Dementia | UW ADRC | 3 | 3 |
| 1 | TZR13 | HCF/High ADNC | 92 | Male | NA | 5.57 | 3_4 | No dementia | ACT | 3 | 3 |
| 1 | TZR14 | HCF/Low ADNC | 88 | Female | NA | 3.5 | 3_3 | No dementia | ACT | 2 | 0 |
| 2 | TZR15 | Sporadic ADD | 85 | Female | NA | 3.52 | 3_3 | Dementia | UW ADRC | 3 | 2 |
| 2 | TZR16 | Autosomal Dominant ADD | 45 | Female | PSEN1 | 6 | 3_3 | Dementia | UW ADRC | 3 | 3 |
| 2 | TZR17 | Sporadic ADD | 62 | Male | NA | 3.92 | 3_4 | Dementia | UW ADRC | 3 | 3 |
| 2 | TZR18 | Autosomal Dominant ADD | 40 | Male | PSEN1 | 15 | NA | Dementia | DIAN | 3 | 3 |
| 2 | TZR19 | HCF/High ADNC | 92 | Female | NA | 4.08 | 3_3 | No dementia | ACT | 2 | 3 |
| 2 | TZR20 | HCF/High ADNC | 87 | Male | NA | 4.5 | 3_3 | No dementia | ACT | 3 | 3 |
| 2 | TZR21 | Sporadic ADD | 86 | Male | NA | 2.67 | 3_4 | Dementia | UW ADRC | 3 | 3 |
| 2 | TZR22 | Sporadic ADD | 91 | Female | NA | 5.75 | 3_4 | Dementia | UW ADRC | 3 | 3 |
| 2 | TZR23 | HCF/High ADNC | 92 | Female | NA | 6 | 4_4 | No dementia | ACT | 2 | 2 |
| 2 | TZR24 | Autosomal Dominant ADD | 60 | Male | PSEN2 | 20.5 | 3_3 | Dementia | UW ADRC | 3 | 3 |
| 2 | TZR25 | HCF/Low ADNC | 94 | Female | NA | 4.85 | 3_3 | No dementia | ACT | 2 | 0 |
| 2 | TZR26 | HCF/Low ADNC | 89 | Male | NA | 5 | 2_3 | No dementia | ACT | 1 | 0 |
| 2 | TZR27 | Sporadic ADD | 91 | Male | NA | 7 | 3_4 | Dementia | ACT | 3 | 2 |
| 2 | TZR28 | Autosomal Dominant ADD | 55 | Male | PSEN2 | 3.98 | 3_4 | Dementia | UW ADRC | 3 | 3 |
| 3 | TZR29 | HCF/High ADNC | 86 | Female | NA | 6.5 | 3_4 | No dementia | ACT | 2 | 2 |
| 3 | TZR30 | Sporadic ADD | 59 | Male | NA | 5.08 | 3_4 | Dementia | UW ADRC | 3 | 3 |
| 3 | TZR31 | Autosomal Dominant ADD | 43 | Male | PSEN1 | 20 | NA | Dementia | DIAN | 3 | 3 |
| 3 | TZR32 | Autosomal Dominant ADD | 45 | Male | PSEN1 | 3.6 | 3_3 | Dementia | UW ADRC | 3 | 3 |
| 3 | TZR33 | HCF/High ADNC | 96 | Female | NA | 6.87 | 3_3 | No dementia | ACT | 2 | 3 |
| 3 | TZR34 | HCF/Low ADNC | 84 | Male | NA | 4.5 | 3_4 | No dementia | ACT | 1 | 0 |
| 3 | TZR35 | Autosomal Dominant ADD | 64 | Male | PSEN2 | 19 | 3_3 | Dementia | UW ADRC | 3 | 2 |
| 3 | TZR36 | HCF/Low ADNC | 95 | Female | NA | 3.08 | 3_4 | No dementia | ACT | 2 | 0 |
| 3 | TZR37 | Sporadic ADD | 93 | Female | NA | 6.5 | 3_3 | Dementia | ACT | 3 | 3 |
| 3 | TZR38 | HCF/High ADNC | 89 | Female | NA | 5.08 | 3_3 | No dementia | ACT | 3 | 3 |
| 3 | TZR39 | Autosomal Dominant ADD | 54 | Female | APP | 6 | 4_4 | Dementia | UW ADRC | 3 | 3 |
| 3 | TZR40 | Autosomal Dominant ADD | 37 | Male | PSEN1 | 7 | 3_3 | Dementia | UW ADRC | 3 | 3 |
| 3 | TZR41 | Sporadic ADD | 94 | Male | NA | 3.85 | 3_3 | Dementia | ACT | 3 | 3 |
| 3 | TZR42 | Sporadic ADD | 85 | Male | NA | 3.25 | 3_3 | Dementia | UW ADRC | 3 | 3 |
| 4 | TZR43 | HCF/High ADNC | 98 | Male | NA | 4 | 3_4 | No dementia | ACT | 3 | 2 |
| 4 | TZR44 | Sporadic ADD | 87 | Female | NA | 5.27 | 3_3 | Dementia | UW ADRC | 3 | 3 |
| 4 | TZR45 | Autosomal Dominant ADD | 39 | Male | PSEN1 | 18 | NA | Dementia | DIAN | 3 | 3 |
| 4 | TZR46 | HCF/Low ADNC | 79 | Female | NA | 2.5 | 3_3 | No dementia | ACT | 1 | 0 |
| 4 | TZR47 | Autosomal Dominant ADD | 42 | Male | PSEN1 | 24 | 3_3 | Dementia | UW ADRC | 3 | 3 |
| 4 | TZR48 | Sporadic ADD | 89 | Female | NA | 5.23 | 3_3 | Dementia | ACT | 3 | 2 |
| 4 | TZR49 | Sporadic ADD | 67 | Female | NA | 4.75 | 3_3 | Dementia | UW ADRC | 3 | 3 |
| 4 | TZR50 | Autosomal Dominant ADD | 49 | Female | PSEN1 | 6.5 | NA | Dementia | DIAN | 3 | 3 |
| 4 | TZR51 | Sporadic ADD | 100 | Male | NA | 4.33 | 3_3 | Dementia | ACT | 3 | 3 |
| 4 | TZR52 | Autosomal Dominant ADD | 58 | Male | PSEN1 | 24 | 2_3 | Dementia | UW ADRC | 3 | 3 |
| 4 | TZR53 | HCF/Low ADNC | 86 | Male | NA | 3.25 | 3_3 | No dementia | ACT | 2 | 0 |
| 4 | TZR54 | Autosomal Dominant ADD | 51 | Female | PSEN1 | 24 | 2_3 | Dementia | UW ADRC | 3 | 3 |
| 4 | TZR55 | Autosomal Dominant ADD | 55 | Male | PSEN2 | 24 | 3_4 | Dementia | UW ADRC | 3 | 3 |
| 4 | TZR56 | HCF/High ADNC | 79 | Male | NA | 2.5 | 3_3 | No dementia | ACT | 2 | 2 |
| 5 | TZR57 | Autosomal Dominant ADD | 72 | Male | PSEN2 | NA | 3_4 | Dementia | UW ADRC | 3 | 3 |
| 5 | TZR58 | Autosomal Dominant ADD | 63 | Male | PSEN1 | 8 | 3_3 | Dementia | UW ADRC | 3 | 3 |
| 5 | TZR59 | Autosomal Dominant ADD | 35 | Female | PSEN1 | 41.8 | NA | Dementia | DIAN | 3 | 3 |
| 5 | TZR60 | Autosomal Dominant ADD | 61 | Male | PSEN1 | 16 | 3_3 | Dementia | UW ADRC | 3 | 3 |
| 5 | TZR61 | Autosomal Dominant ADD | 51 | Male | PSEN1 | 9 | NA | Dementia | DIAN | 3 | 3 |
| 5 | TZR62 | HCF/Low ADNC | 88 | Female | NA | 3.18 | 2_3 | No dementia | ACT | 1 | 0 |

**Supplementary Table 2. Summary metadata table for the hippocampus brain tissue.**

| **Batch** | **Sample Label** | **Condition** | **Age** | **Sex** | **Mutation Status** | **PMI (hrs)** | ***APOE* Alleles** | **Cognitive Status** | **Study Name** | **Braak Stage** | **CERAD Score** |
| --- | --- | --- | --- | --- | --- | --- | --- | --- | --- | --- | --- |
| 1 | HZR01 | Sporadic ADD | 77 | Male | NA | 5.92 | 3_3 | Dementia | ACT | 3 | 3 |
| 1 | HZR02 | Autosomal Dominant ADD | 61 | Male | PSEN1 | 16 | 3_3 | Dementia | UW ADRC | 3 | 3 |
| 1 | HZR03 | HCF/High ADNC | 98 | Male | NA | 4 | 3_4 | No dementia | ACT | 3 | 2 |
| 1 | HZR04 | Autosomal Dominant ADD | 45 | Male | PSEN1 | 3.6 | 3_3 | Dementia | UW ADRC | 3 | 3 |
| 1 | HZR05 | HCF/Low ADNC | 84 | Male | NA | 4.5 | 3_4 | No dementia | ACT | 1 | 0 |
| 1 | HZR06 | HCF/High ADNC | 92 | Male | NA | 5.57 | 3_4 | No dementia | ACT | 3 | 3 |
| 1 | HZR07 | Sporadic ADD | 100 | Male | NA | 4.33 | 3_3 | Dementia | ACT | 3 | 3 |
| 1 | HZR08 | HCF/Low ADNC | 89 | Male | NA | 5 | 2_3 | No dementia | ACT | 1 | 0 |
| 1 | HZR09 | Sporadic ADD | 62 | Male | NA | 3.92 | 3_4 | Dementia | UW ADRC | 3 | 3 |
| 1 | HZR10 | HCF/High ADNC | 78 | Male | NA | 4 | 3_3 | No dementia | ACT | 2 | 2 |
| 1 | HZR11 | HCF/High ADNC | 96 | Female | NA | 6.87 | 3_3 | No dementia | ACT | 2 | 3 |
| 1 | HZR12 | HCF/Low ADNC | 88 | Female | NA | 3.18 | 2_3 | No dementia | ACT | 1 | 0 |
| 1 | HZR13 | HCF/Low ADNC | 88 | Female | NA | 3.5 | 3_3 | No dementia | ACT | 2 | 0 |
| 1 | HZR14 | Sporadic ADD | 89 | Female | NA | 5.23 | 3_3 | Dementia | ACT | 3 | 2 |
| 2 | HZR15 | Sporadic ADD | 63 | Male | NA | 3.33 | 3_3 | Dementia | UW ADRC | 3 | 3 |
| 2 | HZR16 | HCF/High ADNC | 89 | Female | NA | 2.5 | 3_4 | No dementia | ACT | 3 | 3 |
| 2 | HZR17 | HCF/Low ADNC | 95 | Female | NA | 3.08 | 3_4 | No dementia | ACT | 2 | 0 |
| 2 | HZR18 | HCF/High ADNC | 87 | Male | NA | 4.5 | 3_3 | No dementia | ACT | 3 | 3 |
| 2 | HZR19 | HCF/High ADNC | 93 | Male | NA | 6.83 | 3_3 | No dementia | ACT | 3 | 2 |
| 2 | HZR20 | Sporadic ADD | 59 | Male | NA | 5.08 | 3_4 | Dementia | UW ADRC | 3 | 3 |
| 2 | HZR21 | HCF/Low ADNC | 91 | Male | NA | 5 | 3_3 | No dementia | ACT | 2 | 0 |
| 2 | HZR22 | Sporadic ADD | 93 | Female | NA | 6.5 | 3_3 | Dementia | ACT | 3 | 3 |
| 2 | HZR23 | Sporadic ADD | 87 | Female | NA | 5.27 | 3_3 | Dementia | UW ADRC | 3 | 3 |
| 2 | HZR24 | Sporadic ADD | 76 | Male | NA | 3.92 | 3_4 | Dementia | UW ADRC | 3 | 3 |
| 2 | HZR25 | Sporadic ADD | 88 | Male | NA | 4.38 | 3_3 | Dementia | UW ADRC | 3 | 3 |
| 2 | HZR26 | Sporadic ADD | 85 | Female | NA | 3.52 | 3_3 | Dementia | UW ADRC | 3 | 2 |
| 2 | HZR27 | HCF/Low ADNC | 79 | Female | NA | 2.5 | 3_3 | No dementia | ACT | 1 | 0 |
| 2 | HZR28 | HCF/High ADNC | 86 | Female | NA | 6.5 | 3_4 | No dementia | ACT | 2 | 2 |
| 3 | HZR29 | Sporadic ADD | 86 | Male | NA | 2.67 | 3_4 | Dementia | UW ADRC | 3 | 3 |
| 3 | HZR30 | Sporadic ADD | 94 | Male | NA | 3.85 | 3_3 | Dementia | ACT | 3 | 3 |
| 3 | HZR31 | Sporadic ADD | 91 | Female | NA | 5.75 | 3_4 | Dementia | UW ADRC | 3 | 3 |
| 3 | HZR32 | HCF/High ADNC | 92 | Female | NA | 6 | 4_4 | No dementia | ACT | 2 | 2 |
| 3 | HZR33 | Sporadic ADD | 78 | Female | NA | 2.35 | 4_4 | Dementia | UW ADRC | 3 | 2 |
| 3 | HZR34 | HCF/Low ADNC | 73 | Male | NA | 4.5 | 3_3 | No dementia | ACT | 1 | 0 |
| 3 | HZR35 | Sporadic ADD | 67 | Female | NA | 4.75 | 3_3 | Dementia | UW ADRC | 3 | 3 |
| 3 | HZR36 | Sporadic ADD | 85 | Male | NA | 3.25 | 3_3 | Dementia | UW ADRC | 3 | 3 |
| 3 | HZR37 | Sporadic ADD | 90 | Female | NA | 3.85 | 3_3 | Dementia | ACT | 3 | 3 |
| 3 | HZR38 | Autosomal Dominant ADD | 54 | Female | APP | 6 | 4_4 | Dementia | UW ADRC | 3 | 3 |
| 3 | HZR39 | HCF/High ADNC | 89 | Female | NA | 5.08 | 3_3 | No dementia | ACT | 3 | 3 |
| 3 | HZR40 | HCF/Low ADNC | 94 | Female | NA | 4.85 | 3_3 | No dementia | ACT | 2 | 0 |
| 3 | HZR41 | Sporadic ADD | 91 | Male | NA | 7 | 3_4 | Dementia | ACT | 3 | 2 |
| 3 | HZR42 | HCF/High ADNC | 92 | Female | NA | 4.08 | 3_3 | No dementia | ACT | 2 | 3 |
| 3 | HZR43 | Sporadic ADD | 61 | Male | NA | 3 | 4_4 | Dementia | UW ADRC | 3 | 3 |
| 3 | HZR44 | HCF/Low ADNC | 86 | Male | NA | 3.25 | 3_3 | No dementia | ACT | 2 | 0 |

**Supplementary Table 3. Summary metadata table for the inferior parietal lobe (IPL) brain tissue.**

| **Batch** | **Sample Label** | **Condition** | **Age** | **Sex** | **Mutation Status** | **PMI (hrs)** | ***APOE* Alleles** | **Cognitive Status** | **Study Name** | **Braak Stage** | **CERAD Score** |
| --- | --- | --- | --- | --- | --- | --- | --- | --- | --- | --- | --- |
| 1 | PZR01 | HCF/High ADNC | 98 | Male | NA | 4 | 3_4 | No dementia | ACT | 3 | 2 |
| 1 | PZR02 | Autosomal Dominant ADD | 51 | Female | PSEN1 | 24 | 2_3 | Dementia | UW ADRC | 3 | 3 |
| 1 | PZR03 | HCF/Low ADNC | 95 | Female | NA | 3.08 | 3_4 | No dementia | ACT | 2 | 0 |
| 1 | PZR04 | Sporadic ADD | 63 | Male | NA | 3.33 | 3_3 | Dementia | UW ADRC | 3 | 3 |
| 1 | PZR05 | Sporadic ADD | 87 | Female | NA | 5.27 | 3_3 | Dementia | UW ADRC | 3 | 3 |
| 1 | PZR06 | Autosomal Dominant ADD | 55 | Male | PSEN2 | 3.98 | 3_4 | Dementia | UW ADRC | 3 | 3 |
| 1 | PZR07 | Sporadic ADD | 77 | Male | NA | 5.92 | 3_3 | Dementia | ACT | 3 | 3 |
| 1 | PZR08 | Autosomal Dominant ADD | 63 | Male | PSEN1 | 8 | 3_3 | Dementia | UW ADRC | 3 | 3 |
| 1 | PZR09 | Sporadic ADD | 62 | Male | NA | 3.92 | 3_4 | Dementia | UW ADRC | 3 | 3 |
| 1 | PZR10 | HCF/High ADNC | 89 | Female | NA | 2.5 | 3_4 | No dementia | ACT | 3 | 3 |
| 1 | PZR11 | Autosomal Dominant ADD | 43 | Male | PSEN1 | 20 | NA | Dementia | DIAN | 3 | 3 |
| 1 | PZR12 | Sporadic ADD | 61 | Male | NA | 3 | 4_4 | Dementia | UW ADRC | 3 | 3 |
| 1 | PZR13 | HCF/Low ADNC | 89 | Male | NA | 5 | 2_3 | No dementia | ACT | 1 | 0 |
| 1 | PZR14 | HCF/High ADNC | 92 | Female | NA | 4.08 | 3_3 | No dementia | ACT | 2 | 3 |
| 2 | PZR15 | Sporadic ADD | 67 | Female | NA | 4.75 | 3_3 | Dementia | UW ADRC | 3 | 3 |
| 2 | PZR16 | HCF/High ADNC | 89 | Female | NA | 5.08 | 3_3 | No dementia | ACT | 3 | 3 |
| 2 | PZR17 | HCF/High ADNC | 93 | Male | NA | 6.83 | 3_3 | No dementia | ACT | 3 | 2 |
| 2 | PZR18 | Sporadic ADD | 91 | Female | NA | 5.75 | 3_4 | Dementia | UW ADRC | 3 | 3 |
| 2 | PZR19 | Sporadic ADD | 59 | Male | NA | 5.08 | 3_4 | Dementia | UW ADRC | 3 | 3 |
| 2 | PZR20 | HCF/Low ADNC | 86 | Male | NA | 3.25 | 3_3 | No dementia | ACT | 2 | 0 |
| 2 | PZR21 | Autosomal Dominant ADD | 44 | Female | PSEN1 | 17.5 | 3_3 | Dementia | UW ADRC | 3 | 3 |
| 2 | PZR22 | HCF/High ADNC | 92 | Female | NA | 6 | 4_4 | No dementia | ACT | 2 | 2 |
| 2 | PZR23 | Autosomal Dominant ADD | 35 | Female | PSEN1 | 41.8 | NA | Dementia | DIAN | 3 | 3 |
| 2 | PZR24 | Autosomal Dominant ADD | 37 | Male | PSEN1 | 7 | 3_3 | Dementia | UW ADRC | 3 | 3 |
| 2 | PZR25 | Sporadic ADD | 93 | Female | NA | 6.5 | 3_3 | Dementia | ACT | 3 | 3 |
| 2 | PZR26 | HCF/Low ADNC | 94 | Female | NA | 4.85 | 3_3 | No dementia | ACT | 2 | 0 |
| 2 | PZR27 | Autosomal Dominant ADD | 54 | Female | APP | 6 | 4_4 | Dementia | UW ADRC | 3 | 3 |
| 2 | PZR28 | Autosomal Dominant ADD | 45 | Male | PSEN1 | 3.6 | 3_3 | Dementia | UW ADRC | 3 | 3 |
| 3 | PZR29 | Sporadic ADD | 91 | Male | NA | 7 | 3_4 | Dementia | ACT | 3 | 2 |
| 3 | PZR30 | HCF/Low ADNC | 88 | Female | NA | 3.18 | 2_3 | No dementia | ACT | 1 | 0 |
| 3 | PZR31 | Sporadic ADD | 85 | Male | NA | 3.25 | 3_3 | Dementia | UW ADRC | 3 | 3 |
| 3 | PZR32 | Autosomal Dominant ADD | 49 | Female | PSEN1 | 6.5 | NA | Dementia | DIAN | 3 | 3 |
| 3 | PZR33 | Autosomal Dominant ADD | 55 | Male | PSEN2 | 24 | 3_4 | Dementia | UW ADRC | 3 | 3 |
| 3 | PZR34 | HCF/High ADNC | 96 | Female | NA | 6.87 | 3_3 | No dementia | ACT | 2 | 3 |
| 3 | PZR35 | Sporadic ADD | 100 | Male | NA | 4.33 | 3_3 | Dementia | ACT | 3 | 3 |
| 3 | PZR36 | HCF/Low ADNC | 84 | Male | NA | 4.5 | 3_4 | No dementia | ACT | 1 | 0 |
| 3 | PZR37 | HCF/High ADNC | 92 | Male | NA | 5.57 | 3_4 | No dementia | ACT | 3 | 3 |
| 3 | PZR38 | HCF/High ADNC | 78 | Male | NA | 4 | 3_3 | No dementia | ACT | 2 | 2 |
| 3 | PZR39 | Sporadic ADD | 76 | Male | NA | 3.92 | 3_4 | Dementia | UW ADRC | 3 | 3 |
| 3 | PZR40 | Autosomal Dominant ADD | 45 | Female | PSEN1 | 6 | 3_3 | Dementia | UW ADRC | 3 | 3 |
| 3 | PZR41 | Autosomal Dominant ADD | 60 | Male | PSEN2 | 20.5 | 3_3 | Dementia | UW ADRC | 3 | 3 |
| 3 | PZR42 | Sporadic ADD | 78 | Female | NA | 2.35 | 4_4 | Dementia | UW ADRC | 3 | 2 |
| 4 | PZR43 | Sporadic ADD | 86 | Male | NA | 2.67 | 3_4 | Dementia | UW ADRC | 3 | 3 |
| 4 | PZR44 | Sporadic ADD | 94 | Male | NA | 3.85 | 3_3 | Dementia | ACT | 3 | 3 |
| 4 | PZR45 | Autosomal Dominant ADD | 39 | Male | PSEN1 | 18 | NA | Dementia | DIAN | 3 | 3 |
| 4 | PZR46 | HCF/Low ADNC | 88 | Female | NA | 3.5 | 3_3 | No dementia | ACT | 2 | 0 |
| 4 | PZR47 | Sporadic ADD | 85 | Female | NA | 3.52 | 3_3 | Dementia | UW ADRC | 3 | 2 |
| 4 | PZR48 | HCF/High ADNC | 79 | Male | NA | 2.5 | 3_3 | No dementia | ACT | 2 | 2 |
| 4 | PZR49 | HCF/High ADNC | 86 | Female | NA | 6.5 | 3_4 | No dementia | ACT | 2 | 2 |
| 4 | PZR50 | Autosomal Dominant ADD | 51 | Male | PSEN1 | 9 | NA | Dementia | DIAN | 3 | 3 |
| 4 | PZR51 | HCF/High ADNC | 87 | Male | NA | 4.5 | 3_3 | No dementia | ACT | 3 | 3 |
| 4 | PZR52 | Autosomal Dominant ADD | 52 | Female | PSEN1 | NA | 3_4 | Dementia | UW ADRC | 3 | 3 |
| 4 | PZR53 | Autosomal Dominant ADD | 72 | Male | PSEN2 | NA | 3_4 | Dementia | UW ADRC | 3 | 3 |
| 4 | PZR54 | HCF/Low ADNC | 91 | Male | NA | 5 | 3_3 | No dementia | ACT | 2 | 0 |
| 4 | PZR55 | Autosomal Dominant ADD | 61 | Male | PSEN1 | 16 | 3_3 | Dementia | UW ADRC | 3 | 3 |
| 4 | PZR56 | Sporadic ADD | 89 | Female | NA | 5.23 | 3_3 | Dementia | ACT | 3 | 2 |
| 5 | PZR57 | Autosomal Dominant ADD | 64 | Male | PSEN2 | 19 | 3_3 | Dementia | UW ADRC | 3 | 2 |
| 5 | PZR58 | Autosomal Dominant ADD | 39 | Female | PSEN1 | 25 | NA | Dementia | DIAN | 3 | 3 |
| 5 | PZR59 | Autosomal Dominant ADD | 58 | Male | PSEN1 | 24 | 2_3 | Dementia | UW ADRC | 3 | 3 |
| 5 | PZR60 | Autosomal Dominant ADD | 42 | Male | PSEN1 | 24 | 3_3 | Dementia | UW ADRC | 3 | 3 |
| 5 | PZR61 | Autosomal Dominant ADD | 40 | Male | PSEN1 | 15 | NA | Dementia | DIAN | 3 | 3 |

**Supplementary Table 4. Summary metadata table for the caudate nucleus.**

| **Batch** | **Sample Label** | **Condition** | **Age** | **Sex** | **Mutation Status** | **PMI (hrs)** | ***APOE* Alleles** | **Cognitive Status** | **Study Name** | **Braak Stage** | **CERAD Score** |
| --- | --- | --- | --- | --- | --- | --- | --- | --- | --- | --- | --- |
| 1 | CZR01 | Sporadic ADD | 94 | Male | NA | 3.85 | 3_3 | Dementia | ACT | 3 | 3 |
| 1 | CZR02 | HCF/High ADNC | 92 | Male | NA | 5.57 | 3_4 | No dementia | ACT | 3 | 3 |
| 1 | CZR03 | HCF/High ADNC | 87 | Male | NA | 4.5 | 3_3 | No dementia | ACT | 3 | 3 |
| 1 | CZR04 | Autosomal Dominant ADD | 61 | Male | PSEN1 | 16 | 3_3 | Dementia | UW ADRC | 3 | 3 |
| 1 | CZR05 | Autosomal Dominant ADD | 39 | Male | PSEN1 | 18 | NA | Dementia | DIAN | 3 | 3 |
| 1 | CZR06 | Sporadic ADD | 59 | Male | NA | 5.08 | 3_4 | Dementia | UW ADRC | 3 | 3 |
| 1 | CZR07 | Autosomal Dominant ADD | 54 | Female | APP | 6 | 4_4 | Dementia | UW ADRC | 3 | 3 |
| 1 | CZR08 | HCF/Low ADNC | 88 | Female | NA | 3.5 | 3_3 | No dementia | ACT | 2 | 0 |
| 1 | CZR09 | Autosomal Dominant ADD | 42 | Male | PSEN1 | 24 | 3_3 | Dementia | UW ADRC | 3 | 3 |
| 1 | CZR10 | Autosomal Dominant ADD | 63 | Male | PSEN1 | 8 | 3_3 | Dementia | UW ADRC | 3 | 3 |
| 1 | CZR11 | Sporadic ADD | 78 | Female | NA | 2.35 | 4_4 | Dementia | UW ADRC | 3 | 2 |
| 1 | CZR12 | HCF/High ADNC | 92 | Female | NA | 4.08 | 3_3 | No dementia | ACT | 2 | 3 |
| 1 | CZR13 | Autosomal Dominant ADD | 49 | Female | PSEN1 | 6.5 | NA | Dementia | DIAN | 3 | 3 |
| 1 | CZR14 | HCF/Low ADNC | 78 | Male | NA | 4 | 2_3 | No dementia | ACT | 0 | 0 |
| 2 | CZR15 | HCF/Low ADNC | 88 | Female | NA | 3.18 | 2_3 | No dementia | ACT | 1 | 0 |
| 2 | CZR16 | HCF/High ADNC | 92 | Female | NA | 6 | 4_4 | No dementia | ACT | 2 | 2 |
| 2 | CZR17 | Autosomal Dominant ADD | 51 | Male | PSEN1 | 9 | NA | Dementia | DIAN | 3 | 3 |
| 2 | CZR18 | HCF/Low ADNC | 89 | Male | NA | 5 | 2_3 | No dementia | ACT | 1 | 0 |
| 2 | CZR19 | Sporadic ADD | 91 | Male | NA | 7 | 3_4 | Dementia | ACT | 3 | 2 |
| 2 | CZR20 | Autosomal Dominant ADD | 60 | Male | PSEN2 | 20.5 | 3_3 | Dementia | UW ADRC | 3 | 3 |
| 2 | CZR21 | Sporadic ADD | 91 | Female | NA | 5.75 | 3_4 | Dementia | UW ADRC | 3 | 3 |
| 2 | CZR22 | Sporadic ADD | 87 | Female | NA | 5.27 | 3_3 | Dementia | UW ADRC | 3 | 3 |
| 2 | CZR23 | HCF/High ADNC | 96 | Female | NA | 6.87 | 3_3 | No dementia | ACT | 2 | 3 |
| 2 | CZR24 | Autosomal Dominant ADD | 52 | Female | PSEN1 | NA | 3_4 | Dementia | UW ADRC | 3 | 3 |
| 2 | CZR25 | Sporadic ADD | 85 | Male | NA | 3.25 | 3_3 | Dementia | UW ADRC | 3 | 3 |
| 2 | CZR26 | Autosomal Dominant ADD | 45 | Male | PSEN1 | 3.6 | 3_3 | Dementia | UW ADRC | 3 | 3 |
| 2 | CZR27 | Sporadic ADD | 86 | Male | NA | 2.67 | 3_4 | Dementia | UW ADRC | 3 | 3 |
| 2 | CZR28 | HCF/High ADNC | 93 | Male | NA | 6.83 | 3_3 | No dementia | ACT | 3 | 2 |
| 3 | CZR29 | Sporadic ADD | 76 | Male | NA | 3.92 | 3_4 | Dementia | UW ADRC | 3 | 3 |
| 3 | CZR30 | HCF/Low ADNC | 95 | Female | NA | 3.08 | 3_4 | No dementia | ACT | 2 | 0 |
| 3 | CZR31 | Sporadic ADD | 67 | Female | NA | 4.75 | 3_3 | Dementia | UW ADRC | 3 | 3 |
| 3 | CZR32 | Autosomal Dominant ADD | 78 | Female | PSEN2 | 7 | 2_3 | Dementia | UW ADRC | 3 | 3 |
| 3 | CZR33 | Autosomal Dominant ADD | 37 | Male | PSEN1 | 7 | 3_3 | Dementia | UW ADRC | 3 | 3 |
| 3 | CZR34 | HCF/High ADNC | 86 | Female | NA | 6.5 | 3_4 | No dementia | ACT | 2 | 2 |
| 3 | CZR35 | Sporadic ADD | 77 | Male | NA | 5.92 | 3_3 | Dementia | ACT | 3 | 3 |
| 3 | CZR36 | HCF/High ADNC | 98 | Male | NA | 4 | 3_4 | No dementia | ACT | 3 | 2 |
| 3 | CZR37 | Sporadic ADD | 89 | Female | NA | 5.23 | 3_3 | Dementia | ACT | 3 | 2 |
| 3 | CZR38 | Sporadic ADD | 90 | Female | NA | 3.85 | 3_3 | Dementia | ACT | 3 | 3 |
| 3 | CZR39 | HCF/Low ADNC | 86 | Male | NA | 3.25 | 3_3 | No dementia | ACT | 2 | 0 |
| 3 | CZR40 | Autosomal Dominant ADD | 58 | Male | PSEN1 | 24 | 2_3 | Dementia | UW ADRC | 3 | 3 |
| 3 | CZR41 | HCF/High ADNC | 89 | Female | NA | 5.08 | 3_3 | No dementia | ACT | 3 | 3 |
| 3 | CZR42 | Autosomal Dominant ADD | 39 | Female | PSEN1 | 25 | NA | Dementia | DIAN | 3 | 3 |
| 4 | CZR43 | Autosomal Dominant ADD | 45 | Female | PSEN1 | 6 | 3_3 | Dementia | UW ADRC | 3 | 3 |
| 4 | CZR44 | HCF/High ADNC | 79 | Male | NA | 2.5 | 3_3 | No dementia | ACT | 2 | 2 |
| 4 | CZR45 | HCF/Low ADNC | 79 | Female | NA | 2.5 | 3_3 | No dementia | ACT | 1 | 0 |
| 4 | CZR46 | Autosomal Dominant ADD | 55 | Male | PSEN2 | 3.98 | 3_4 | Dementia | UW ADRC | 3 | 3 |
| 4 | CZR47 | Sporadic ADD | 63 | Male | NA | 3.33 | 3_3 | Dementia | UW ADRC | 3 | 3 |
| 4 | CZR48 | HCF/Low ADNC | 73 | Male | NA | 4.5 | 3_3 | No dementia | ACT | 1 | 0 |
| 4 | CZR49 | Sporadic ADD | 93 | Female | NA | 6.5 | 3_3 | Dementia | ACT | 3 | 3 |
| 4 | CZR50 | HCF/Low ADNC | 91 | Male | NA | 5 | 3_3 | No dementia | ACT | 2 | 0 |
| 4 | CZR51 | Sporadic ADD | 88 | Male | NA | 4.38 | 3_3 | Dementia | UW ADRC | 3 | 3 |
| 4 | CZR52 | Autosomal Dominant ADD | 43 | Male | PSEN1 | 20 | NA | Dementia | DIAN | 3 | 3 |
| 4 | CZR53 | Autosomal Dominant ADD | 55 | Male | PSEN2 | 24 | 3_4 | Dementia | UW ADRC | 3 | 3 |
| 4 | CZR54 | Sporadic ADD | 85 | Female | NA | 3.52 | 3_3 | Dementia | UW ADRC | 3 | 2 |
| 4 | CZR55 | Autosomal Dominant ADD | 64 | Male | PSEN2 | 19 | 3_3 | Dementia | UW ADRC | 3 | 2 |
| 4 | CZR56 | Autosomal Dominant ADD | 35 | Female | PSEN1 | 41.8 | NA | Dementia | DIAN | 3 | 3 |
| 4 | CZR57 | Sporadic ADD | 61 | Male | NA | 3 | 4_4 | Dementia | UW ADRC | 3 | 3 |
| 4 | CZR58 | HCF/Low ADNC | 84 | Male | NA | 4.5 | 3_4 | No dementia | ACT | 1 | 0 |

**Supplementary Table 5. Mass spectrometry run order table for superior and middle temporal gyri (SMTG).**

| **Run Order** | **Sample Name** | **Sample type** |
| --- | --- | --- |
| 1 | QC01 | System Suitability |
| 2 | QC02 | System Suitability |
| 3 | QC03 | System Suitability |
| 4 | QC04 | System Suitability |
| 5 | Equil-01 | Equilibration |
| 6 | Equil-02 | Equilibration |
| 7 | Equil-03 | Equilibration |
| 8 | Equil-04 | Equilibration |
| 9 | Equil-05 | Equilibration |
| 10 | Equil-06 | Equilibration |
| 11 | Equil-07 | Equilibration |
| 12 | Equil-08 | Equilibration |
| 13 | Equil-09 | Equilibration |
| 14 | TZR03 | Batch1 Sample Quant |
| 15 | TZR08 | Batch1 Sample Quant |
| 16 | TZR07 | Batch1 Sample Quant |
| 17 | TZR04 | Batch1 Sample Quant |
| 18 | TZR09 | Batch1 Sample Quant |
| 19 | TZR10 | Batch1 Sample Quant |
| 20 | TZR01 | Batch1 Sample Quant |
| 21 | TZR12 | Batch1 Sample Quant |
| 22 | QC06 | System Suitability |
| 23 | Batch 1 Pool | B1 Chr library 400-500 |
| 24 | Batch 1 Pool | B1 Chr library 500-600 |
| 25 | Batch 1 Pool | B1 Chr library 600-700 |
| 26 | Batch 1 Pool | B1 Chr library 700-800 |
| 27 | Batch 1 Pool | B1 Chr library 800-900 |
| 28 | Batch 1 Pool | B1 Chr library 900-1000 |
| 29 | QC07 | System Suitability |
| 30 | HADT01 | Batch QC (Brain QC) |
| 31 | TZR13 | Batch1 Sample Quant |
| 32 | TZR06 | Batch1 Sample Quant |
| 33 | TZR11 | Batch1 Sample Quant |
| 34 | TRPR01 | Batch Reference (Batch1 pool) |
| 35 | TZR02 | Batch1 Sample Quant |
| 36 | TZR05 | Batch1 Sample Quant |
| 37 | TZR14 | Batch1 Sample Quant |
| 38 | QC08 | System Suitability |
| 39 | TZR26 | Batch 2 Sample Quant |
| 40 | TZR21 | Batch 2 Sample Quant |
| 41 | TZR20 | Batch 2 Sample Quant |
| 42 | TZR27 | Batch 2 Sample Quant |
| 43 | TZR24 | Batch 2 Sample Quant |
| 44 | TZR16 | Batch 2 Sample Quant |
| 45 | TZR22 | Batch 2 Sample Quant |
| 46 | TZR17 | Batch 2 Sample Quant |
| 47 | QC09 | System Suitability |
| 48 | Batch 2 Pool | B2 Chr library 400-500 |
| 49 | Batch 2 Pool | B2 Chr library 500-600 |
| 50 | Batch 2 Pool | B2 Chr library 600-700 |
| 51 | Batch 2 Pool | B2 Chr library 700-800 |
| 52 | Batch 2 Pool | B2 Chr library 800-900 |
| 53 | Batch 2 Pool | B2 Chr library 900-1000 |
| 54 | QC10 | System Suitability |
| 55 | TZR15 | Batch 2 Sample Quant |
| 56 | TZR25 | Batch 2 Sample Quant |
| 57 | TZR18 | Batch 2 Sample Quant |
| 58 | HADT02 | Batch QC (Brain QC) |
| 59 | TZR23 | Batch 2 Sample Quant |
| 60 | TZR19 | Batch 2 Sample Quant |
| 61 | TZR28 | Batch 2 Sample Quant |
| 62 | TRPR02 | Batch Reference (Batch1 pool) |
| 63 | QC11 | System Suitability |
| 64 | TZR35 | Batch 3 Sample Quant |
| 65 | TZR37 | Batch 3 Sample Quant |
| 66 | TZR33 | Batch 3 Sample Quant |
| 67 | TRPR03 | Batch Reference (Batch1 pool) |
| 68 | TZR39 | Batch 3 Sample Quant |
| 69 | TZR32 | Batch 3 Sample Quant |
| 70 | TZR42 | Batch 3 Sample Quant |
| 71 | HADT03 | Batch QC (Brain QC) |
| 72 | QC12 | System Suitability |
| 73 | Batch 3 Pool | B3 Chr library 400-500 |
| 74 | Batch 3 Pool | B3 Chr library 500-600 |
| 75 | Batch 3 Pool | B3 Chr library 600-700 |
| 76 | Batch 3 Pool | B3 Chr library 700-800 |
| 77 | Batch 3 Pool | B3 Chr library 800-900 |
| 78 | Batch 3 Pool | B3 Chr library 900-1000 |
| 79 | QC13 | System Suitability |
| 80 | TZR30 | Batch 3 Sample Quant |
| 81 | TZR29 | Batch 3 Sample Quant |
| 82 | TZR38 | Batch 3 Sample Quant |
| 83 | TZR40 | Batch 3 Sample Quant |
| 84 | TZR41 | Batch 3 Sample Quant |
| 85 | TZR34 | Batch 3 Sample Quant |
| 86 | TZR36 | Batch 3 Sample Quant |
| 87 | TZR31 | Batch 3 Sample Quant |
| 88 | QC14 | System Suitability |
| 89 | TRPR05 | Batch Reference (Batch1 pool) |
| 90 | TZR48 | Batch 4 and 5 Sample Quant |
| 91 | TZR57 | Batch 4 and 5 Sample Quant |
| 92 | HADTP05 | Batch QC (Brain QC) |
| 93 | TZR46 | Batch 4 and 5 Sample Quant |
| 94 | TZR50 | Batch 4 and 5 Sample Quant |
| 95 | TZR47 | Batch 4 and 5 Sample Quant |
| 96 | TRPR04 | Batch Reference (Batch1 pool) |
| 97 | QC15 | System Suitability |
| 98 | HADT04 | Batch QC (Brain QC) |
| 99 | TZR51 | Batch 4 and 5 Sample Quant |
| 100 | TZR55 | Batch 4 and 5 Sample Quant |
| 101 | TZR45 | Batch 4 and 5 Sample Quant |
| 102 | TZR61 | Batch 4 and 5 Sample Quant |
| 103 | TZR44 | Batch 4 and 5 Sample Quant |
| 104 | TZR54 | Batch 4 and 5 Sample Quant |
| 105 | TZR43 | Batch 4 and 5 Sample Quant |
| 106 | QC16 | System Suitability |
| 107 | Batch 4 and 5 Pool | B4-B5 Chr library 400-500 |
| 108 | Batch 4 and 5 Pool | B4-B5 Chr library 500-600 |
| 109 | Batch 4 and 5 Pool | B4-B5 Chr library 600-700 |
| 110 | Batch 4 and 5 Pool | B4-B5 Chr library 700-800 |
| 111 | Batch 4 and 5 Pool | B4-B5 Chr library 800-900 |
| 112 | Batch 4 and 5 Pool | B4-B5 Chr library 900-1000 |
| 113 | QC17 | System Suitability |
| 114 | TZR49 | Batch 4 and 5 Sample Quant |
| 115 | TZR58 | Batch 4 and 5 Sample Quant |
| 116 | TZR59 | Batch 4 and 5 Sample Quant |
| 117 | TZR53 | Batch 4 and 5 Sample Quant |
| 118 | TZR60 | Batch 4 and 5 Sample Quant |
| 119 | TZR56 | Batch 4 and 5 Sample Quant |
| 120 | TZR52 | Batch 4 and 5 Sample Quant |
| 121 | TZR62 | Batch 4 and 5 Sample Quant |
| 122 | QC18 | System Suitability |

**Supplementary Table 6. Mass spectrometry run order table for the hippocampus.**

| **Run Order** | **Sample Name** | **Sample type** |
| --- | --- | --- |
| 1 | QC01 | System Suitability |
| 2 | QC02 | System Suitability |
| 3 | QC03 | System Suitability |
| 4 | QC04 | System Suitability |
| 5 | Equil-01 | Equilibration |
| 6 | Equil-02 | Equilibration |
| 7 | Equil-03 | Equilibration |
| 8 | Equil-04 | Equilibration |
| 9 | Equil-05 | Equilibration |
| 10 | Equil-06 | Equilibration |
| 11 | Equil-07 | Equilibration |
| 12 | Equil-08 | Equilibration |
| 13 | Equil-09 | Equilibration |
| 14 | QC05 | System Suitability |
| 15 | HZR11 | Batch1 Sample Quant |
| 16 | HRPR01 | Batch Reference (Batch1 pool) |
| 17 | HZR10 | Batch1 Sample Quant |
| 18 | HZR07 | Batch1 Sample Quant |
| 19 | HZR01 | Batch1 Sample Quant |
| 20 | HAD01 | Batch QC (Brain QC) |
| 21 | HZR02 | Batch1 Sample Quant |
| 22 | HZR08 | Batch1 Sample Quant |
| 23 | QC06 | System Suitability |
| 24 | Batch 1 Pool | B1 Chr library 400-500 |
| 25 | Batch 1 Pool | B1 Chr library 500-600 |
| 26 | Batch 1 Pool | B1 Chr library 600-700 |
| 27 | Batch 1 Pool | B1 Chr library 700-800 |
| 28 | Batch 1 Pool | B1 Chr library 800-900 |
| 29 | Batch 1 Pool | B1 Chr library 900-1000 |
| 30 | QC07 | System Suitability |
| 31 | HZR12 | Batch1 Sample Quant |
| 32 | HZR13 | Batch1 Sample Quant |
| 33 | HZR04 | Batch1 Sample Quant |
| 34 | HZR06 | Batch1 Sample Quant |
| 35 | HZR09 | Batch1 Sample Quant |
| 36 | HZR05 | Batch1 Sample Quant |
| 37 | HZR14 | Batch1 Sample Quant |
| 38 | HZR03 | Batch1 Sample Quant |
| 39 | QC08 | System Suitability |
| 40 | HZR15 | Batch 2 Sample Quant |
| 41 | HZR20 | Batch 2 Sample Quant |
| 42 | HZR24 | Batch 2 Sample Quant |
| 43 | HZR16 | Batch 2 Sample Quant |
| 44 | HZR22 | Batch 2 Sample Quant |
| 45 | HZR28 | Batch 2 Sample Quant |
| 46 | HZR25 | Batch 2 Sample Quant |
| 47 | HZR18 | Batch 2 Sample Quant |
| 48 | QC09 | System Suitability |
| 49 | Batch 2 Pool | B2 Chr library 400-500 |
| 50 | Batch 2 Pool | B2 Chr library 500-600 |
| 51 | Batch 2 Pool | B2 Chr library 600-700 |
| 52 | Batch 2 Pool | B2 Chr library 700-800 |
| 53 | Batch 2 Pool | B2 Chr library 800-900 |
| 54 | Batch 2 Pool | B2 Chr library 900-1000 |
| 55 | QC10 | System Suitability |
| 56 | HRPR02 | Batch Reference (Batch1 pool) |
| 57 | HZR17 | Batch 2 Sample Quant |
| 58 | HZR23 | Batch 2 Sample Quant |
| 59 | HAD02 | Batch QC (Brain QC) |
| 60 | HZR26 | Batch 2 Sample Quant |
| 61 | HZR21 | Batch 2 Sample Quant |
| 62 | HZR19 | Batch 2 Sample Quant |
| 63 | HZR27 | Batch 2 Sample Quant |
| 64 | QC11 | System Suitability |
| 65 | HZR34 | Batch 3 Sample Quant |
| 66 | HZR40 | Batch 3 Sample Quant |
| 67 | HZR44 | Batch 3 Sample Quant |
| 68 | HZR31 | Batch 3 Sample Quant |
| 69 | HZR32 | Batch 3 Sample Quant |
| 70 | HZR35 | Batch 3 Sample Quant |
| 71 | HZR41 | Batch 3 Sample Quant |
| 72 | QC12 | System Suitability |
| 73 | Batch 3 Pool | B3 Chr library 400-500 |
| 74 | Batch 3 Pool | B3 Chr library 500-600 |
| 75 | Batch 3 Pool | B3 Chr library 600-700 |
| 76 | Batch 3 Pool | B3 Chr library 700-800 |
| 77 | Batch 3 Pool | B3 Chr library 800-900 |
| 78 | Batch 3 Pool | B3 Chr library 900-1000 |
| 79 | QC13 | System Suitability |
| 80 | HZR29 | Batch 3 Sample Quant |
| 81 | HRPR03b | Batch Reference (Batch1 pool) |
| 82 | HZR37 | Batch 3 Sample Quant |
| 83 | HAD03a | Batch QC (Brain QC) |
| 84 | HZR38 | Batch 3 Sample Quant |
| 85 | HZR43 | Batch 3 Sample Quant |
| 86 | HZR42 | Batch 3 Sample Quant |
| 87 | QC14 | System Suitability |
| 88 | HZR30 | Batch 3 Sample Quant |
| 89 | HAD03b | Batch QC (Brain QC) |
| 90 | HZR36 | Batch 3 Sample Quant |
| 91 | HRPR03a | Batch Reference (Batch1 pool) |
| 92 | HZR39 | Batch 3 Sample Quant |
| 93 | HZR33 | Batch 3 Sample Quant |
| 94 | QC15 | System Suitability |

**Supplementary Table 7. Mass spectrometry run order table for the inferior parietal lobule (IPL).**

| **Run Order** | **Sample Name** | **Sample type** |
| --- | --- | --- |
| 1 | QC01 | System Suitability |
| 2 | QC02 | System Suitability |
| 3 | QC03 | System Suitability |
| 4 | QC04 | System Suitability |
| 5 | Equil-01 | Equilibration |
| 6 | Equil-02 | Equilibration |
| 7 | Equil-03 | Equilibration |
| 8 | Equil-04 | Equilibration |
| 9 | Equil-05 | Equilibration |
| 10 | Equil-06 | Equilibration |
| 11 | Equil-07 | Equilibration |
| 12 | Equil-08 | Equilibration |
| 13 | QC05 | System Suitability |
| 14 | PZR05 | Batch1 Sample Quant |
| 15 | PZR10 | Batch1 Sample Quant |
| 16 | PZR01 | Batch1 Sample Quant |
| 17 | PZR14 | Batch1 Sample Quant |
| 18 | PZR09 | Batch1 Sample Quant |
| 19 | PZR13 | Batch1 Sample Quant |
| 20 | PZR04 | Batch1 Sample Quant |
| 21 | PZR12 | Batch1 Sample Quant |
| 22 | QC06 | System Suitability |
| 23 | Batch 1 Pool | B1 Chr library 400-500 |
| 24 | Batch 1 Pool | B1 Chr library 500-600 |
| 25 | Batch 1 Pool | B1 Chr library 600-700 |
| 26 | Batch 1 Pool | B1 Chr library 700-800 |
| 27 | Batch 1 Pool | B1 Chr library 800-900 |
| 28 | Batch 1 Pool | B1 Chr library 900-1000 |
| 29 | QC07 | System Suitability |
| 30 | PZR11 | Batch1 Sample Quant |
| 31 | PZR03 | Batch1 Sample Quant |
| 32 | PZR06 | Batch1 Sample Quant |
| 33 | PZR08 | Batch1 Sample Quant |
| 34 | HADP01 | Batch QC (Brain QC) |
| 35 | PZR02 | Batch1 Sample Quant |
| 36 | PRPR01 | Batch Reference (Batch1 pool) |
| 37 | PZR07 | Batch1 Sample Quant |
| 38 | QC08 | System Suitability |
| 39 | PZR17 | Batch 2 Sample Quant |
| 40 | PZR21 | Batch 2 Sample Quant |
| 41 | PZR23 | Batch 2 Sample Quant |
| 42 | PZR27 | Batch 2 Sample Quant |
| 43 | PZR22 | Batch 2 Sample Quant |
| 44 | PZR26 | Batch 2 Sample Quant |
| 45 | PZR20 | Batch 2 Sample Quant |
| 46 | PZR19 | Batch 2 Sample Quant |
| 47 | QC09 | System Suitability |
| 48 | Batch 2 Pool | B2 Chr library 400-500 |
| 49 | Batch 2 Pool | B2 Chr library 500-600 |
| 50 | Batch 2 Pool | B2 Chr library 600-700 |
| 51 | Batch 2 Pool | B2 Chr library 700-800 |
| 52 | Batch 2 Pool | B2 Chr library 800-900 |
| 53 | Batch 2 Pool | B2 Chr library 900-1000 |
| 54 | QC10 | System Suitability |
| 55 | PZR25 | Batch 2 Sample Quant |
| 56 | PZR28 | Batch 2 Sample Quant |
| 57 | PZR16 | Batch 2 Sample Quant |
| 58 | PZR24 | Batch 2 Sample Quant |
| 59 | PZR18 | Batch 2 Sample Quant |
| 60 | PRPR02 | Batch Reference (Batch1 pool) |
| 61 | HADP02 | Batch QC (Brain QC) |
| 62 | PZR15 | Batch 2 Sample Quant |
| 63 | QC11 | System Suitability |
| 64 | PZR39 | Batch 3 Sample Quant |
| 65 | PZR34 | Batch 3 Sample Quant |
| 66 | PZR31 | Batch 3 Sample Quant |
| 67 | PZR42 | Batch 3 Sample Quant |
| 68 | PZR33 | Batch 3 Sample Quant |
| 69 | PRPR03 | Batch Reference (Batch1 pool) |
| 70 | PZR30 | Batch 3 Sample Quant |
| 71 | PZR35 | Batch 3 Sample Quant |
| 72 | QC12 | System Suitability |
| 73 | Batch 3 Pool | B3 Chr library 400-500 |
| 74 | Batch 3 Pool | B3 Chr library 500-600 |
| 75 | Batch 3 Pool | B3 Chr library 600-700 |
| 76 | Batch 3 Pool | B3 Chr library 700-800 |
| 77 | Batch 3 Pool | B3 Chr library 800-900 |
| 78 | Batch 3 Pool | B3 Chr library 900-1000 |
| 79 | QC13 | System Suitability |
| 80 | HADP03 | Batch QC (Brain QC) |
| 81 | PZR41 | Batch 3 Sample Quant |
| 82 | PZR32 | Batch 3 Sample Quant |
| 83 | PZR40 | Batch 3 Sample Quant |
| 84 | PZR29 | Batch 3 Sample Quant |
| 85 | PZR36 | Batch 3 Sample Quant |
| 86 | PZR37 | Batch 3 Sample Quant |
| 87 | PZR38 | Batch 3 Sample Quant |
| 88 | QC14 | System Suitability |
| 89 | PZR57 | Batch 4 and 5 Sample Quant |
| 90 | PZR48 | Batch 4 and 5 Sample Quant |
| 91 | PRPR04 | Batch Reference (Batch1 pool) |
| 92 | PRPR05 | Batch Reference (Batch1 pool) |
| 93 | PZR49 | Batch 4 and 5 Sample Quant |
| 94 | PZR58 | Batch 4 and 5 Sample Quant |
| 95 | PZR43 | Batch 4 and 5 Sample Quant |
| 96 | PZR45 | Batch 4 and 5 Sample Quant |
| 97 | QC15 | System Suitability |
| 98 | Batch 4 and 5 Pool | B4-B5 Chr library 400-500 |
| 99 | Batch 4 and 5 Pool | B4-B5 Chr library 500-600 |
| 100 | Batch 4 and 5 Pool | B4-B5 Chr library 600-700 |
| 101 | Batch 4 and 5 Pool | B4-B5 Chr library 700-800 |
| 102 | Batch 4 and 5 Pool | B4-B5 Chr library 800-900 |
| 103 | Batch 4 and 5 Pool | B4-B5 Chr library 900-1000 |
| 104 | QC16 | System Suitability |
| 105 | PZR55 | Batch 4 and 5 Sample Quant |
| 106 | PZR51 | Batch 4 and 5 Sample Quant |
| 107 | PZR61 | Batch 4 and 5 Sample Quant |
| 108 | HADP05 | Batch QC (Brain QC) |
| 109 | PZR60 | Batch 4 and 5 Sample Quant |
| 110 | PZR50 | Batch 4 and 5 Sample Quant |
| 111 | PZR52 | Batch 4 and 5 Sample Quant |
| 112 | PZR47 | Batch 4 and 5 Sample Quant |
| 113 | QC17 | System Suitability |
| 114 | PZR44 | Batch 4 and 5 Sample Quant |
| 115 | PZR59 | Batch 4 and 5 Sample Quant |
| 116 | HADP04 | Batch QC (Brain QC) |
| 117 | PZR54 | Batch 4 and 5 Sample Quant |
| 118 | PZR53 | Batch 4 and 5 Sample Quant |
| 119 | PZR46 | Batch 4 and 5 Sample Quant |
| 120 | PZR56 | Batch 4 and 5 Sample Quant |
| 121 | QC18 | System Suitability |

**Supplementary Table 8. Mass spectrometry run order table for the caudate nucleus.**

| **Run Order** | **Sample Name** | **Sample type** |
| --- | --- | --- |
| 1 | QC01 | System Suitability |
| 2 | QC02 | System Suitability |
| 3 | QC03 | System Suitability |
| 4 | QC04 | System Suitability |
| 5 | QC05 | System Suitability |
| 6 | QC06 | System Suitability |
| 7 | QC07 | System Suitability |
| 8 | QC08 | System Suitability |
| 9 | QC09 | System Suitability |
| 10 | QC10 | System Suitability |
| 12 | QC11 | System Suitability - new vial |
| 13 | QC12 | System Suitability |
| 14 | Equil-01 | Equilibration |
| 15 | Equil-02 | Equilibration |
| 16 | Equil-03 | Equilibration |
| 17 | Equil-04 | Equilibration |
| 18 | Equil-05 | Equilibration |
| 19 | Equil-06 | Equilibration |
| 20 | Equil-07 | Equilibration |
| 21 | Equil-08 | Equilibration |
| 22 | QC13 | System Suitability |
| 23 | CZR07 | Batch1 Sample Quant |
| 24 | CZR08 | Batch1 Sample Quant |
| 25 | CZR10 | Batch1 Sample Quant |
| 26 | CZR13 | Batch1 Sample Quant |
| 27 | CRPR01 | Batch Reference (Batch1 pool) |
| 28 | CZR05 | Batch1 Sample Quant |
| 29 | CZR03 | Batch1 Sample Quant |
| 30 | CZR09 | Batch1 Sample Quant |
| 31 | QC14 | System Suitability |
| 32 | Batch 1 Pool | B1 Chr library 400-500 |
| 33 | Batch 1 Pool | B1 Chr library 500-600 |
| 34 | Batch 1 Pool | B1 Chr library 600-700 |
| 35 | Batch 1 Pool | B1 Chr library 700-800 |
| 36 | Batch 1 Pool | B1 Chr library 800-900 |
| 37 | Batch 1 Pool | B1 Chr library 900-1000 |
| 38 | QC15 | System Suitability |
| 39 | CZR01 | Batch1 Sample Quant |
| 40 | CZR12 | Batch1 Sample Quant |
| 41 | CZR06 | Batch1 Sample Quant |
| 42 | CZR14 | Batch1 Sample Quant |
| 43 | CZR02 | Batch1 Sample Quant |
| 44 | HADC01 | Batch QC (Brain QC) |
| 45 | CZR11 | Batch1 Sample Quant |
| 46 | CZR04 | Batch1 Sample Quant |
| 47 | QC16 | System Suitability |
| 48 | CZR19 | Batch 2 Sample Quant |
| 49 | CZR24 | Batch 2 Sample Quant |
| 50 | CZR28 | Batch 2 Sample Quant |
| 51 | CZR21 | Batch 2 Sample Quant |
| 52 | CZR23 | Batch 2 Sample Quant |
| 53 | CZR27 | Batch 2 Sample Quant |
| 54 | CZR16 | Batch 2 Sample Quant |
| 55 | CZR25 | Batch 2 Sample Quant |
| 56 | QC17 | System Suitability |
| 57 | Batch 2 Pool | B2 Chr library 400-500 |
| 58 | Batch 2 Pool | B2 Chr library 500-600 |
| 59 | Batch 2 Pool | B2 Chr library 600-700 |
| 60 | Batch 2 Pool | B2 Chr library 700-800 |
| 61 | Batch 2 Pool | B2 Chr library 800-900 |
| 62 | Batch 2 Pool | B2 Chr library 900-1000 |
| 63 | QC18 | System Suitability |
| 64 | CZR20 | Batch 2 Sample Quant |
| 65 | HADC02 | Batch QC (Brain QC) |
| 66 | CZR15 | Batch 2 Sample Quant |
| 67 | CZR22 | Batch 2 Sample Quant |
| 68 | CZR17 | Batch 2 Sample Quant |
| 69 | CZR18 | Batch 2 Sample Quant |
| 70 | CZR26 | Batch 2 Sample Quant |
| 71 | CRPR02 | Batch Reference (Batch1 pool) |
| 72 | QC19 | System Suitability |
| 73 | CZR31 | Batch 3 Sample Quant |
| 74 | CZR33 | Batch 3 Sample Quant |
| 75 | CZR37 | Batch 3 Sample Quant |
| 76 | CZR36 | Batch 3 Sample Quant |
| 77 | CZR42 | Batch 3 Sample Quant |
| 78 | CZR41 | Batch 3 Sample Quant |
| 79 | CZR38 | Batch 3 Sample Quant |
| 80 | CZR40 | Batch 3 Sample Quant |
| 81 | QC20 | System Suitability |
| 82 | Batch 3 Pool | B3 Chr library 400-500 |
| 83 | Batch 3 Pool | B3 Chr library 500-600 |
| 84 | Batch 3 Pool | B3 Chr library 600-700 |
| 85 | Batch 3 Pool | B3 Chr library 700-800 |
| 86 | Batch 3 Pool | B3 Chr library 800-900 |
| 87 | Batch 3 Pool | B3 Chr library 900-1000 |
| 88 | QC21 | System Suitability |
| 89 | CZR32 | Batch 3 Sample Quant |
| 90 | CZR30 | Batch 3 Sample Quant |
| 91 | CZR35 | Batch 3 Sample Quant |
| 92 | CZR34 | Batch 3 Sample Quant |
| 93 | CZR29 | Batch 3 Sample Quant |
| 94 | HADC03 | Batch QC (Brain QC) |
| 95 | CRPR03 | Batch Reference (Batch1 pool) |
| 96 | CZR39 | Batch 3 Sample Quant |
| 97 | QC22 | System Suitability |
| 98 | CZR55 | Batch 4 Sample Quant |
| 99 | CRPR04a | Batch 4 Sample Quant |
| 100 | CRPR04b | Batch Reference (Batch1 pool) |
| 101 | CZR49 | Batch Reference (Batch1 pool) |
| 102 | CZR51 | Batch 4 Sample Quant |
| 103 | HADC04a | Batch QC (Brain QC) |
| 104 | CZR46 | Batch 4 Sample Quant |
| 105 | QC23 | System Suitability |
| 106 | Batch 4 Pool | B4 Chr library 400-500 |
| 107 | Batch 4 Pool | B4 Chr library 500-600 |
| 108 | Batch 4 Pool | B4 Chr library 600-700 |
| 109 | Batch 4 Pool | B4 Chr library 700-800 |
| 110 | Batch 4 Pool | B4 Chr library 800-900 |
| 111 | Batch 4 Pool | B4 Chr library 900-1000 |
| 112 | QC24 | System Suitability |
| 113 | CZR57 | Batch 4 Sample Quant |
| 114 | CZR47 | Batch 4 Sample Quant |
| 115 | CZR53 | Batch 4 Sample Quant |
| 116 | CZR48 | Batch 4 Sample Quant |
| 117 | CZR44 | Batch 4 Sample Quant |
| 118 | CZR43 | Batch 4 Sample Quant |
| 119 | HADC04b | Batch QC (Brain QC) |
| 120 | QC25 | System Suitability |
| 121 | CZR56 | Batch 4 Sample Quant |
| 122 | CZR45 | Batch 4 Sample Quant |
| 123 | CZR58 | Batch 4 Sample Quant |
| 124 | CZR54 | Batch 4 Sample Quant |
| 125 | CZR50 | Batch 4 Sample Quant |
| 126 | CZR52 | Batch 4 Sample Quant |
| 127 | QC26 | System Suitability |
